# Visual input into the *Drosophila melanogaster* mushroom body

**DOI:** 10.1101/2020.02.07.935924

**Authors:** Jinzhi Li, Brennan Dale Mahoney, Miles Solomon Jacob, Sophie Jeanne Cécile Caron

## Abstract

The ability to integrate input from different sensory systems is a fundamental property of many brains. Yet, the patterns of neuronal connectivity that underlie such multisensory integration remain poorly characterized. The *Drosophila melanogaster* mushroom body — an associative center required for the formation of olfactory and visual memories — is an ideal system to investigate how different sensory channels converge in higher-order brain centers. The neurons connecting the mushroom body to the olfactory system have been described in great detail, but input from other sensory systems remains poorly defined. Here, we use a range of anatomical and genetic techniques to identify two novel types of mushroom body input neuron that connect visual processing centers — namely the lobula and the posterior lateral protocerebrum — to the dorsal accessory calyx of the mushroom body. Together with previous work that described a pathway conveying visual information from the medulla to the ventral accessory calyx of the mushroom body (Vogt et al., 2016), our study defines a second, parallel pathway that is anatomically poised to convey information from the visual system to the dorsal accessory calyx. This connectivity pattern — the segregation of the visual information into two separate pathways — could be a fundamental feature of the neuronal architecture underlying multisensory integration in associative brain centers.

## INTRODUCTION

Sensory systems use different strategies to detect specific physical features of the outside world. For instance, the olfactory system contains many different types of sensory neuron that are each specialized in detecting a specific class of volatile chemicals. Through only two neuronal layers, olfactory information — the identity of an odor and its concentration — is relayed to higher brain centers (Leinwand and Chalasani, 2011). In contrast, the visual system contains far fewer types of sensory neuron but, through numerous neuronal layers, it relays a range of highly processed information — for instance, color, brightness, motion and shape — to higher brain centers (Baden et al., 2020). Thus, higher brain centers have to integrate different types of processed information, bind that information into a coherent representation of the outside world, and use such representations to guide behavior (Yau et al., 2015). How higher brain centers achieve this feat remains largely unknown. This gap in our knowledge mainly stems from the fact that higher brain centers are formed by a large number of neurons, and that the projection neurons conveying information from different sensory systems to these centers often remain poorly characterized. This makes it difficult to understand whether there are specific patterns of neuronal connectivity that enable multisensory integration and what the nature of these patterns are. Deciphering the fundamental neuronal mechanisms that underlie multisensory integration requires a model system such as the *Drosophila melanogaster* mushroom body, which consists of a relatively small number of neurons whose connections can be charted reliably.

The *Drosophila* mushroom body is formed by ~2,000 neurons — called the Kenyon cells — and has long been studied for its essential role in the formation of olfactory associative memories (Aso et al., 2014; Hige, 2018). The identity of the projection neurons that connect the olfactory system to the mushroom body — and the way Kenyon cells integrate input from these neurons — has been characterized in great detail, highlighting fundamental connectivity patterns that enable this higher brain center to represent olfactory information efficiently (Bates et al., 2020; Caron et al., 2013; Tanaka et al., 2012a, 2012b; Zheng et al., 2018). Evidence in *Drosophila melanogaster* shows that the mushroom body is more than an olfactory center, as it is also required for the formation of visual and gustatory associative memories (Liu et al., 1999; Masek and Scott, 2010; Vogt et al., 2014). However, the identity of the neurons that connect the mushroom body to other sensory systems remains poorly characterized. Thus, a first step towards understanding how the mushroom body integrates multisensory information is to identify such non-olfactory mushroom body input neurons and the genetic tools necessary to manipulate these neurons.

The mushroom body receives its input through its calyx and sends its output through its lobes. The calyx — a morphologically distinct neuropil containing the synapses formed between projection neurons and Kenyon cells — can be divided into four, non-overlapping regions: one main calyx as well as three accessory calyces named the dorsal, lateral and ventral accessory calyces (Aso et al., 2014; Yagi et al., 2016). The five output lobes — the α, α’, β, β’ and γ lobes — contain the synapses formed between Kenyon cells, mushroom body output neurons and dopaminergic neurons (Aso et al., 2014). With respect to these input and output regions, Kenyon cells can be divided into seven distinct types (Aso et al., 2014). Of these seven types, five types — the α/β_c_, α/β_s_, α’/β’_ap_, α’/β’_m_ and γ_main_ Kenyon cells — extend their dendrites only into the main calyx and their axons along one or two lobes. Most of the neurons that project to the main calyx emerge from the antennal lobe, the primary olfactory center in the *Drosophila* brain. Thus α/β_c_, α/β_s_, α’/β’_ap_, α’/β’_m_ and α_main_ Kenyon cells receive input primarily from the olfactory system (Caron et al., 2013; Aso et al., 2014; Zheng et al., 2018).

In contrast, the two other classes of Kenyon cells do not extend their dendrites into the main calyx. Instead, the α/β_p_ Kenyon cells extend their dendrites into the dorsal accessory calyx — avoiding completely the main, lateral and ventral accessory calyces — and their axons along the α and β lobes. Likewise, the γ_d_ Kenyon cells extend their dendrites exclusively into the ventral accessory calyx and their axons along the γ lobe (Aso et al., 2014; Vogt et al., 2016). Thus, both the α/β_p_ and γ_d_ Kenyon cells are anatomically poised to receive non-olfactory input. There is evidence suggesting that the ventral accessory calyx receives input from the medulla, a region of the optic lobe that specializes in processing brightness and color (Morante and Desplan, 2008; Vogt et al., 2016). Furthermore, a recent study suggests that the dorsal accessory calyx is a multisensory center that integrates input from multiple sensory pathways including the olfactory, gustatory and visual systems (Yagi et al., 2016).

Here, we report a strategy that uses a combination of genetic tools — including transgenic lines that drive expression in few neurons and a photo-labelling technique used to identify individual neurons and their presynaptic partners — to characterize the input neurons of the α/β_p_ Kenyon cells. We identify two novel types of mushroom body input neuron that, together, form about half of the total input the α/β_p_ Kenyon cells receive in the dorsal accessory calyx. The first neuronal type — henceforth referred to as _LO_PNs — consists of a neuron that projects from the lobula, a region of the optic lobe specialized in detecting visual features such as shape and motion. The second type of neuron — henceforth referred to as _PLP_PNs — consists of projection neurons that emerge from the posterior lateral protocerebrum, a brain region that receives input from the optic lobe (Otsuna and Ito, 2006; Keleş and Frye, 2017; Wu et al., 2016). Interestingly, _LO_PN and _PLP_PNs do not project to the ventral accessory calyx and do not connect to the γ_d_ Kenyon cells. Based on these findings, we conclude that there are two parallel pathways that convey visual information to the mushroom body: a pathway projecting from the medulla to the γ_d_ Kenyon cells and another pathway projecting from the lobula and posterior lateral protocerebrum to the α/β_p_ Kenyon cells.

## RESULTS

### Neurons projecting to the dorsal accessory calyx emerge from different brain regions

The dorsal accessory calyx is a neuropil formed from the synapses connecting ~90 α/β_p_ Kenyon cells to their input neurons (Figure 1A; (Aso et al., 2014)). Using transgenic lines that drive expression specifically in the α/β_p_ Kenyon cells (the *R85D07-GAL4_DBD_, RI3FO2-GAL4_AD_* transgenic line also known as the MB371-GAL4) and transgenic lines that express a photo-activatable form of GFP or PA-GFP (a combination of the *UAS-C3PA-GFP* and *UAS-SPA-GFP* transgenic lines, henceforth referred to as *UAS-PA-GFP),* we photo-labelled individual α/β_p_ Kenyon cells similarly as described in a previous study (Aso et al., 2014; Datta et al., 2008; Ruta et al., 2010). We found that individual α/β_p_ Kenyon cells extend on average 5 ± 1 claw-shaped dendritic terminals (*n* = 13) exclusively into the dorsal accessory calyx (Figure 1B, D) and project their axons along the α and β lobes (not depicted). The overall morphology of α/β_p_ Kenyon cells is similar to the morphology of other types of α/β Kenyon cell; for instance, individual α/β_s_ Kenyon cells extend on average 6 ± 1 claw-shaped dendritic terminals (*n* = 12) exclusively in the main calyx (Figure 1C) and project their axons along the α and β lobes (not depicted). It is worth noting that the claws formed by the α/β_p_ Kenyon cells are much smaller in diameter (1.8 ± 0.4 μm; *n* = 9 claws) than the claws formed by the α/β_s_ Kenyon cells (3.0 ± 0.4 μm; *n* = 10 claws; Figure 1B-C (white arrows), E). It is also worth noting that the border of the dorsal accessory calyx is looser than the compact and well-defined circular border of the main calyx. Individual α/β_p_ Kenyon cells can extend dendrites further away from the core of the dorsal accessory calyx, resulting in an irregularly shaped calyx (Figure 1D). Thus, the α/β_p_ Kenyon cells resemble α/β_s_ and α/β_c_ Kenyon cells but also differ from them in one major way: the dendrites of the α/β_s_ and α/β_c_ Kenyon cells exclusively innervate the main calyx — a region known to receive most of its input from the olfactory system — whereas dendrites of the α/β_p_ Kenyon cells exclusively innervate the dorsal accessory calyx — a poorly characterized region of the mushroom body.

**Figure 1.**
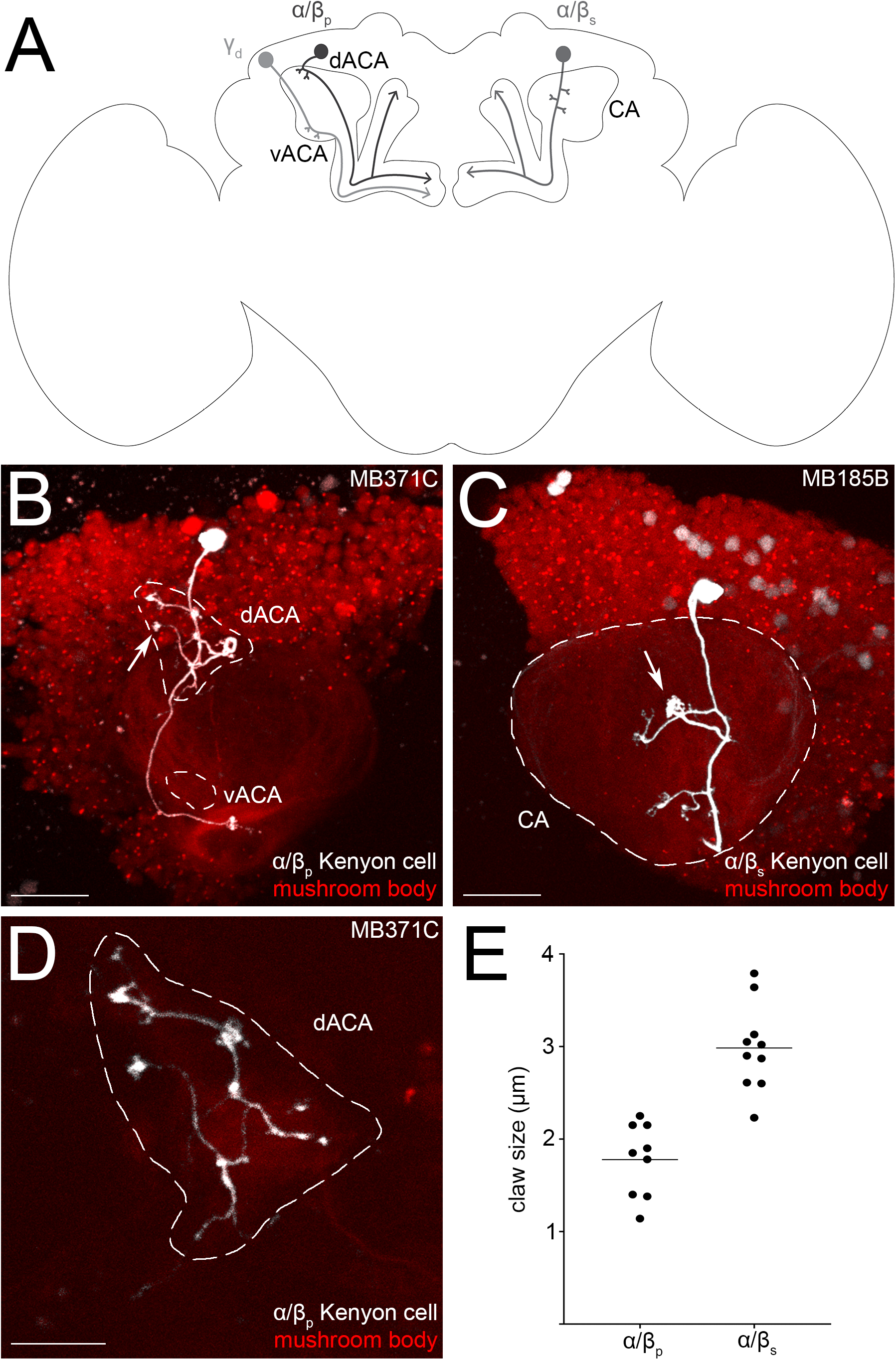
The dorsal accessory calyx of the *Drosophila melanogaster* mushroom body as defined by the claw-shaped dendrites of the α/β_p_ Kenyon cells. (A) A schematic of the *Drosophila* brain shows three types of Kenyon cell associated with the different mushroom body calyces: the α/β_p_ Kenyon cells (dark grey, left) extend their claw-shaped dendrites in the dorsal accessory calyx (dACA), the α/β_s_ (medium grey, right) — as well as the α’/β’_AP_, α’/β’_m_ and γ_main_ Kenyon cells (not depicted) — extend their claw-shaped dendrites in the main calyx (CA) and the γ_d_ Kenyon cells (light grey, left) extend their claw-shaped dendrites in the ventral accessory calyx (vACA). All Kenyon cells extend their axons along one or two lobes (arrowheads). (B-C) A single α/β_p_ (B) or α/β_s_ (C) Kenyon cell (white) of the mushroom body (red) was photo-labeled; (B) the photo-labeled α/β_p_ Kenyon cell extends claw-shaped dendritic terminals (arrows) in the dorsal accessory calyx (dACA, white dashed outline) but not in the ventral accessory calyx (vACA, white dashed outline) or main calyx (not outlined); (C) the photo-labeled α/β_s_ Kenyon cell extends clawshaped dendritic terminals (arrows) in the main calyx (CA, white dashed outline) but not in the dorsal accessory calyx or the ventral accessory calyx (not outlined). (D) The dorsal accessory calyx is loose and irregularly shaped but its core can be visualized in the red channel. (E) α/β_p_ Kenyon cells form smaller claws than the α/β_s_ Kenyon cells. The following genotypes were used in this figure: (B) *yw/yw,MB247-DsRed^unknown^,UAS-C3PA-GFP^unknown^/UAS-SPA-GFP^attP40^;UAS-C3PA-GFP^attP2^, UAS-C3PA-GFP^VK00005^, UAS-C3PA-GFP^VK00027^/MB371C-splitGAL4 [R13F02-GAL4AD^VK00027^, R85D07-GAL4_DBD_^attP2^];* and (C): *yw/yw;MB247-DsRed^unknown^,UAS-C3PA-GFP^unknown^/R52H09-GAL4_AD_^attp40^;UAS-C3PA^attp2^,UAS-C3PA^VK00005^,UAS-C3PA^VK00027^/MB185B-splitGAL4 [R52H09-GAL4_AD_^attP40^, R18F09-GAL4DBD^attP2^]*;. Scale bars are 20 μm (B-C) and 10 μm (D).

To identify neurons that project to the dorsal accessory calyx and connect to the α/β_p_ Kenyon cells, we used a targeted photo-labeling technique that was adapted from previously published techniques (Aso et al., 2014; Datta et al., 2008; Ruta et al., 2010). In order to only photo-label the neurons projecting to the dorsal accessory calyx, and not the α/β_p_ Kenyon cells, we used a combination of transgenes that, in concert, drive the expression of PA-GFP in all neurons except the Kenyon cells (the *N-Synaptobrevin-GAL4, MB247-GAL80* and *UAS-PA-GFP* transgenes); instead, Kenyon cells were labeled with the red fluorescent protein DsRed using the *MB247-DsRed* transgene (Figure 2A). The brains of adult flies carrying all the aforementioned transgenes were dissected and imaged using two-photon microscopy (Figure 2B-C). Guided by the expression of DsRed, we targeted the dorsal accessory calyx — which is clearly distinct from the main calyx — with high-energy light in order to photo-convert PA-GFP specifically in the neurons projecting to that area (Figure 2C, white dashed outline). Upon photo-labeling, the somata of the neurons that express PA-GFP, and that project either dendrites or axons into the dorsal accessory calyx, were labeled (Figure 2D-E). On average, 71 ± 22 neurons (*n* = 22) were photo-labeled. The somata of these neurons are located in seven distinct clusters that are distributed across the brain (Figure 2A, D-E). On the anterior side of the brain, we found three clusters: one located near the antennal lobe (cluster AL containing 3 ± 3 neurons), one located in the optic lobe (cluster OL containing 1 ± 2 neurons) and one located near the anterior ventral lateral protocerebrum (cluster AVLP containing 6 ± 3 neurons) (Figure 2D, F). On the posterior side of the brain, we found four clusters: one located in the superior medial protocerebrum (cluster SMP containing 4 ± 2 neurons), one located in the superior lateral protocerebrum (cluster SLP containing 20 ± 8 neurons), one located in the lateral horn (cluster LH containing 34 ± 12 neurons) and one located in the posterior lateral protocerebrum (cluster PLP containing 3 ± 2 neurons) (Figure 2E-F). Altogether, these results suggest that the dorsal accessory calyx receives input from a diverse and distributed collection of projection neurons that can be divided into seven clusters.

**Figure 2.**
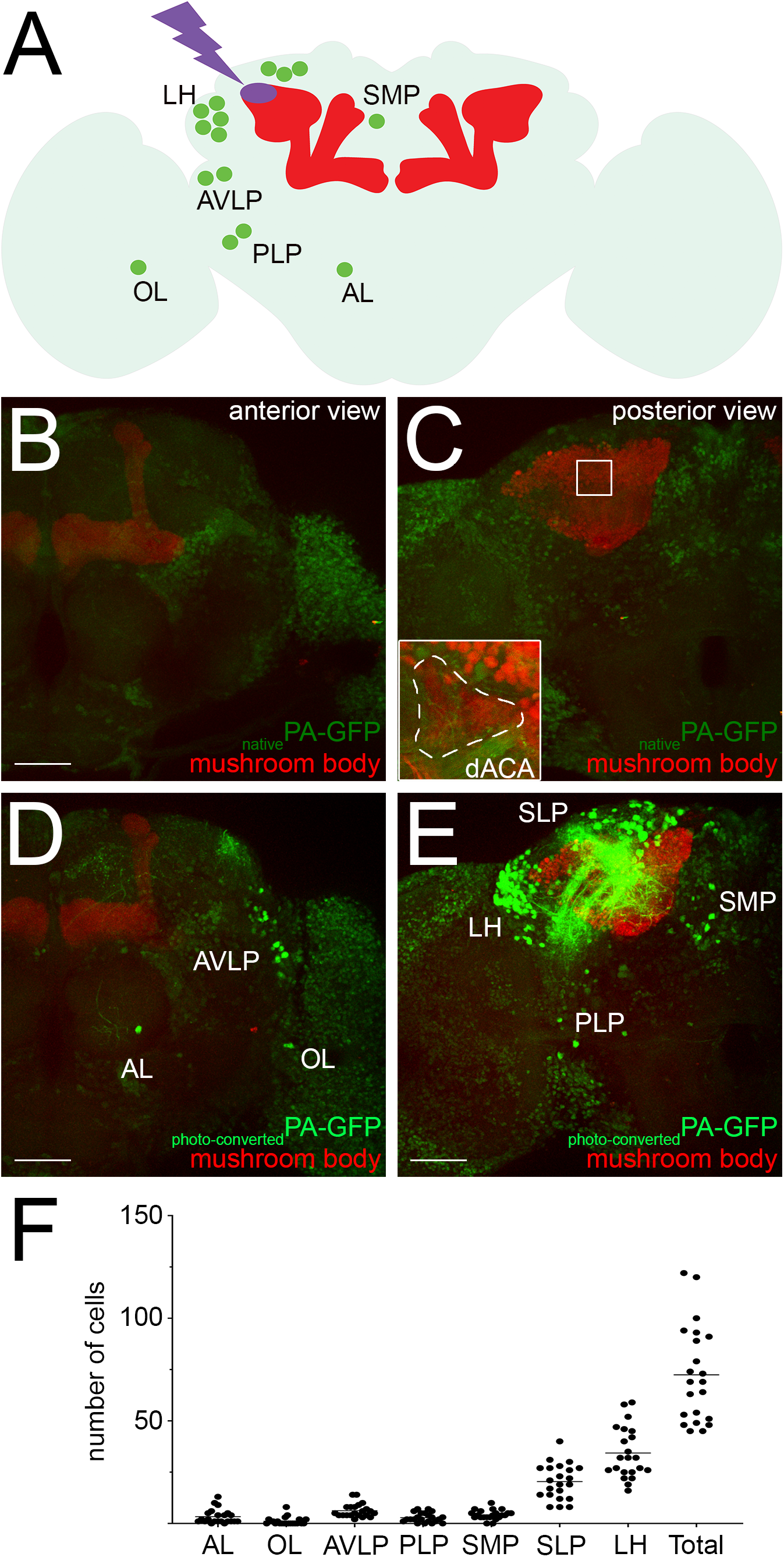
Identification of neurons projecting to the dorsal accessory calyx using *en masse* photo-labeling. (A) A schematic of the *Drosophila* brain shows the seven clusters of neurons identified by photo-labeling the dorsal accessory calyx of the mushroom body. (B-C) All neurons — except for Kenyon cells (red) — express PA-GFP (light green) showing weak fluorescence in structures located on the anterior (B) and posterior (C) sides of the brain. (C) The dorsal accessory calyx (dACA) is innervated by the α/β_p_ Kenyon cells — and no other Kenyon cells — and is clearly visible in the red channel, forming a irregular shape on the dorsal-anterior side of the main calyx; (C, inset) the region outlined by the white dashed line was targeted for photo-labeling. (D-E) Upon photo-activation of PA-GFP in the dorsal accessory calyx, seven clusters of photo-labeled neurons (bright green) are clearly distinguishable in different brain regions located on both the anterior (D) and posterior (E) sides of the brain. The location of these clusters where defined based on the low background fluorescence visible in the unlabeled remaining neurons that were not photo-labeled but that express PA-GFP (light green). (F) This procedure recovered a total of 71 neuronal cell bodies (71 ± 22, *n* = 22) in seven different clusters located near or in the antennal lobe (AL), optic lobe (OL), anterior ventral lateral protocerebrum (AVLP), lateral horn (LH), superior lateral protocerebrum (SLP), superior medial protocerebrum (SMP) and posterior lateral protocerebrum (PLP). The following genotype was used in this figure: *ywlyw;MB247-DsRed^unkown^,UAS-C3PA-GFP^unkown^/MB247-GAL80^unkown^, UAS-SPA-GFP^attP40^; UAS-C3PA-GFP^attP2^, UAS-C3PA-GFP^VK00005^, UAS-C3PA-GFP^VK00027^/N-Synaptobrevin-GAL4^2-1^*;. Scale bars are 50 μm in all panels.

### Identification of transgenic lines driving expression in the dorsal accessory calyx input neurons

While *en masse* photo-labeling of the neurons projecting to the areas within and around the dorsal accessory calyx gives a good approximation of the number and identity of neurons possibly connecting to the α/β_p_ Kenyon cells, this technique cannot confidently identify true pre-synaptic partners. Additionally, the amount of photo-activated PA-GFP within individual neurons varies significantly between trials, and because a large number of neurons project to the dorsal accessory calyx, it is difficult to resolve the morphology of individual neurons using this technique. We, therefore, sought to identify GAL4 transgenic lines that drive expression specifically in these neurons. To identify such lines, we carried out an anatomical screen using the FlyLight collection of GAL4 transgenic lines (Figure 3A; (Jenett et al., 2012)). As a first step, we screened through the FlyLight database — an online catalogue that reports the expression patterns of ~7,000 GAL4 driver lines — and selected 267 transgenic lines which that highlighted neuronal processes in the dorsal accessory calyx (Figure 3B-C).

**Figure 3.**
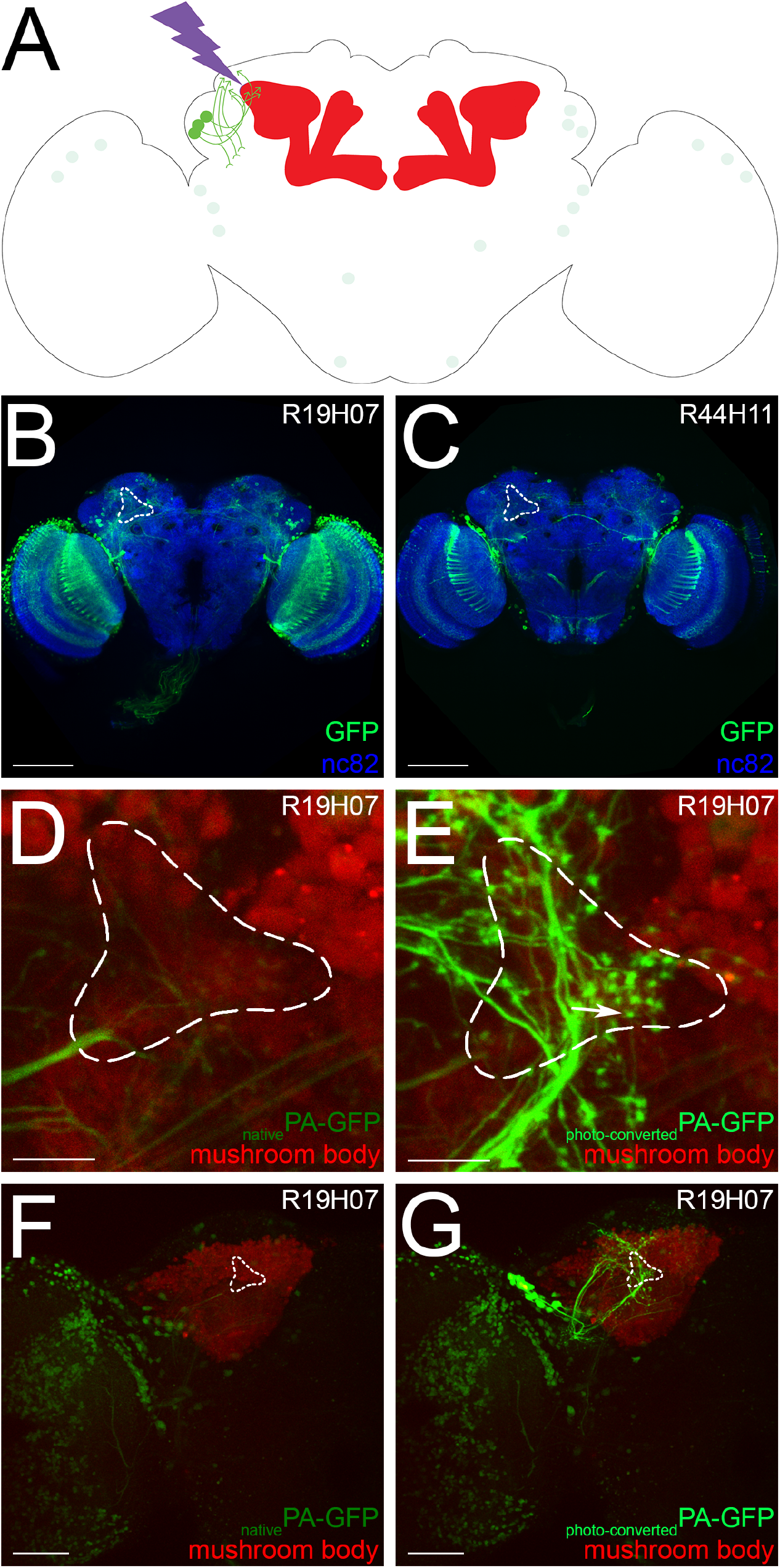
Anatomical screen to identify transgenic lines driving expression in the neurons projecting to the dorsal accessory calyx. (A) A schematic of the *Drosophila* brain shows how transgenic lines driving expression in potential dorsal accessory calyx input neurons were identified by photo-labeling; the schematic shows the identification of a transgenic line driving expression in the _PLP_PNs as an example. (B-C) First, the expression patterns of each of the transgenic lines available in the FlyLight database was inspected: transgenic lines clearly driving expression strongly (B, *R19H07-GAL4)* or weakly (C, *R44H11-GAL4)* in neurons projecting to or from the dorsal accessory calyx (white dashed outline) were selected for further investigation. These images were obtained from the FlyLight website. (D, E) The dorsal accessory calyx (white dashed outline) of the mushroom body (red) was visualized in the selected lines, here *R19H07*-GAL4, and targeted for photo-activation by designing a mask that exposed the outlined region to high energy light; transgenic lines driving expression in a few neurons extending clear axonal terminals in the dorsal accessory calyx (E, arrow) were selected for further investigation. (F, G) Before photo-labeling, weak fluorescence is visible in many different neurons (F) and, upon photolabeling, the neurons projecting to the dorsal accessory calyx (white dashed outline) are visible (G). The following genotype was used in the D-G panels: *yw/yw;MB247-DsRed^unknown^, UAS-C3PA-GFP^unknown^/UAS-C3PA-GFP^attP40^; UAS-C3PA^attP2^, UAS-C3PA^VK00005^, UAS-C3PA^VK00027^/R19H07-GAL4^attP2^*;. Scale bars are 100 μm (B-C), 10 μm (D-E) and 50 μm (F-G).

As a second step, we specifically labeled neurons that project to the dorsal accessory calyx using the same technique as described above but with a different combination of transgenes (the *R_line-GAL4, UAS-PA-GFP* and *MB247-DsRed* transgenes). The brains of adult flies that carry this combination of transgenes were dissected and imaged using two-photon microscopy (Figure 3D, F). Guided by the expression of DsRed, we targeted the dorsal accessory calyx with high-energy light in order to convert PA-GFP specifically in the neurons that project to that region (Figure 3D, white dashed outline). We screen through the 267 pre-selected transgenic lines and identified 10 lines that we chose to investigate further based on two criteria. First, we selected lines that showed a strong signal after photo-labeling and clear pre-synaptic terminals near or in the dorsal accessory calyx (Figure 3E, G; Table 1). Possible pre-synaptic terminals — bouton-shaped (Figure 3E, white arrow) — were distinguished from possible post-synaptic terminals — mesh-shaped — based on the recovered photo-labeling signal. Second, we selected lines based on the strength and specificity of their expression patterns: lines expressing at high level in a few neurons led to clear photolabeling signals, making it easier to characterize these putative input neurons further (Figure 3G).

**Table 1.** Neurons projecting to the dorsal accessory calyx. The neurons identified in the screen were divided into three types — _LO_PN, _PLP_PN and _AL_PN — based on the location of their dendrites — lobula (LO), posterior lateral protocerebrum (PLP) and antennal lobe (AL). The _AL_PN type was further divided into four subtypes — _AL_PN1, _AL_PN2, _AL_PN3 and _AL_PN4 — based on the fact that each one of these neurons were identified using different transgenic lines. The asterisk demarks the only neuron identified by our study that is similar to a neuron described in the Yagi et al. (2016) study.

| Type of neuron | Location of cell body | Number of neurons | Driver line(s) | Connections to $\alpha/\beta_p$ Kenyon cells | |
| --- | --- | --- | --- | --- | --- |
|  |  |  |  | GRASP | Dye/photo-labeling technique |
| **LoPN\***<br>(OCLT1) | PLP cluster | 1 | R44H11 | strong signal | 4.1% ($n = 27$) |
| | | | R72D07 | strong signal | 3.0% ($n = 27$) |
|  |  |  | R95F09 | strong signal | n/A |
| **PLP**PN | LH cluster | 13 | R19H07 | strong signal | All: 14.1% ($n = 24$)<br>Single: 1.9% ($n = 32$) |
|  |  |  | R20G07 | weak signal |  |
| **AL**PN1 | AL cluster | 1 | R30E11 | weak signal | 0 ($n = 26$) |
|  |  |  | R31C03 | weak signal |  |
| **AL**PN2 | AL cluster | 1 | R53B06 | weak signal | 0 ($n = 30$) |
| **AL**PN3 | AL cluster | 1 | R11F08 | no signal | 0 ($n = 26$) |
|  |  |  | R12C04 | no signal |  |
| **AL**PN4 | AL cluster | 1 | R11F07 | no signal | 0 ($n = 25$) |

Finally, we determined whether the neurons identified with these lines are pre-synaptic partners of the α/β_p_ Kenyon cells. To this end, we used an activity-dependent version of the GFP-reconstitution across synaptic partners (GRASP) technique (Figure 4A) (Feinberg et al., 2008; Macpherson et al., 2015). In this technique, the green fluorescent protein (GFP) is split into two complementary fragments — the GFP1-10 and GFP11 fragments — that do not fluoresce when expressed alone; the GFP1-10 fragment is tagged to an anchor that specifically targets the pre-synaptic membrane (syb::spGFP1-10), whereas the GFP11 fragment is tagged to an anchor that targets the membrane (spGFP11::CD4). When both fragments are in close contact — in this case, when the syb::spGFP1-10 fragment is released at an active synapse — GFP molecules can be reconstituted and recover their fluorescence. The reconstitution of GFP molecules, as indicated by the presence of fluorescent GFP speckles, suggests that the neurons expressing the two complementary fragments form functional synapses (Macpherson et al., 2015). In our study, the syb::spGFP1-10 fragment was expressed in the putative input neurons using the different transgenic lines identified in our screen as well as two transgenic lines that a previous study found to drive expression in putative dorsal accessory calyx input neurons (Figure 4B-M, Table 1; (Yagi et al., 2016)). The expression of the spGFP11::CD4 fragment was driven in most of the Kenyon cells using the *MB247-LEXA* transgenic line. We look for GFP speckles in the core of the dorsal accessory calyx, which is clearly visible in the red channel as an irregularly shaped structure anterior to the main calyx (Figure 4B, white dashed outline). We detected a large number of GFP speckles in the core of the dorsal accessory calyx for four of the ten lines that were identified in the screen as well as for the two transgenic lines identified by the Yagi *et al.* study (Figure 4B-E, 4L-M and Table 1). Additionally, we detected a small number of GFP speckles in the same region for three other transgenic lines (Figure 4F-H and Table 1). It is worth noting that some of the GFP speckles lay outside of the red signal; most likely these speckles are formed by Kenyon cells that extend dendrites outside of the core of the dorsal accessory calyx (Figure 4B-H, 4L-M). Finally, we did not detect any GFP speckles for the remaining three lines (Figure 4I-K and Table 1). Altogether, our screen identified at least nine transgenic lines that drive expression in neurons that form synapses with the α/β_p_ Kenyon cells of the dorsal accessory calyx.

**Figure 4.**
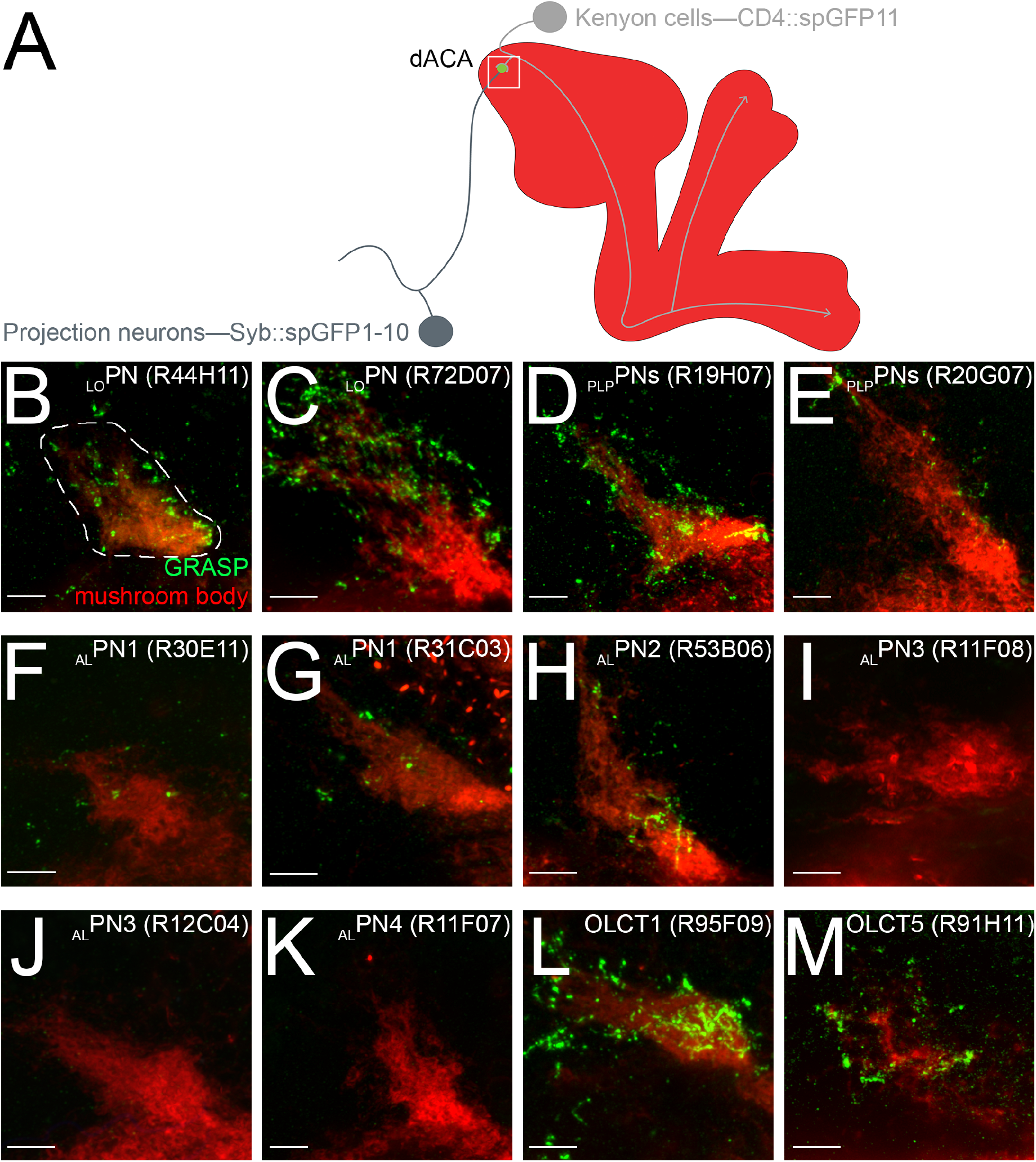
Identification of the transgenic lines driving expression in neurons presynaptic to the α/β_p_ Kenyon cells. (A) A schematic of the *Drosophila* brain shows how the GFP Reconstitution Across Synaptic Partners (GRASP) technique was used to determine whether the selected transgenic lines drive expression in neurons that are presynaptic to the α/β_p_ Kenyon cells. The expression of the spGFP1-10 fragment was driven in the putative input neurons (dark grey) — using the various transgenic lines identified in the screen — and the expression of the spGFP11 fragment was driven in the mushroom body (red) using a transgenic line driving in most Kenyon cells; α/β_p_ Kenyon cell (light grey) can be distinguished from other Kenyon cells because they are the only Kenyon cells projecting to the dorsal accessory calyx. Reconstituted GFP molecules (here depicted by a green circle) are visible in the dorsal accessory calyx of the mushroom body (red) when the spGFP^1-10^ fragment is expressed in synaptic partners. The dorsal accessory calyx was imaged (white square). (B-M) A strong (B-E, L-M) or weak (F-H) GRASP signal was detected in some of the selected transgenic lines; no GRASP signal was detected in other lines (I-K). The following genotypes were used in this figure: *yw/yw;UAS-syb::spGFP1-10^unknown^, LEXAop-CD4::spGFP11^unknown^/LEXAop-tdTomato^attP5^; R_line-GAL4^attP2^* (as indicated on the *panel)/MB247-LEXA^unkown^;.* Scale bars are 5 μm in all panels.

### _LO_PN connects the lobula to the α/β_p_ Kenyon cells

To characterize further the morphology of the projection neurons that connect to the α/β_p_ Kenyon cells, we photo-labeled individual neurons using the transgenic lines recovered from our screen that showed reliable GRASP signal. To this end, neurons were photo-labeled in the dissected brains of flies carrying the *R_line-GAL4* and *UAS-PA-GFP* transgenes; these brains were then immuno-stained and imaged using confocal microscopy. We first focused our attention on the transgenic lines in which we had found the strongest GRASP signal (Figure 4B-E and Table 1). Photo-labeling of neurons that project to the dorsal accessory calyx using the *R44H11-GAL4* and *R72D07-GAL4* transgenic lines revealed a single neuron (Figure 5A). In each line, a neuron projecting from the lobula to the dorsal accessory calyx was clearly photo-labeled (Figure 5B-C).

**Figure 5.**
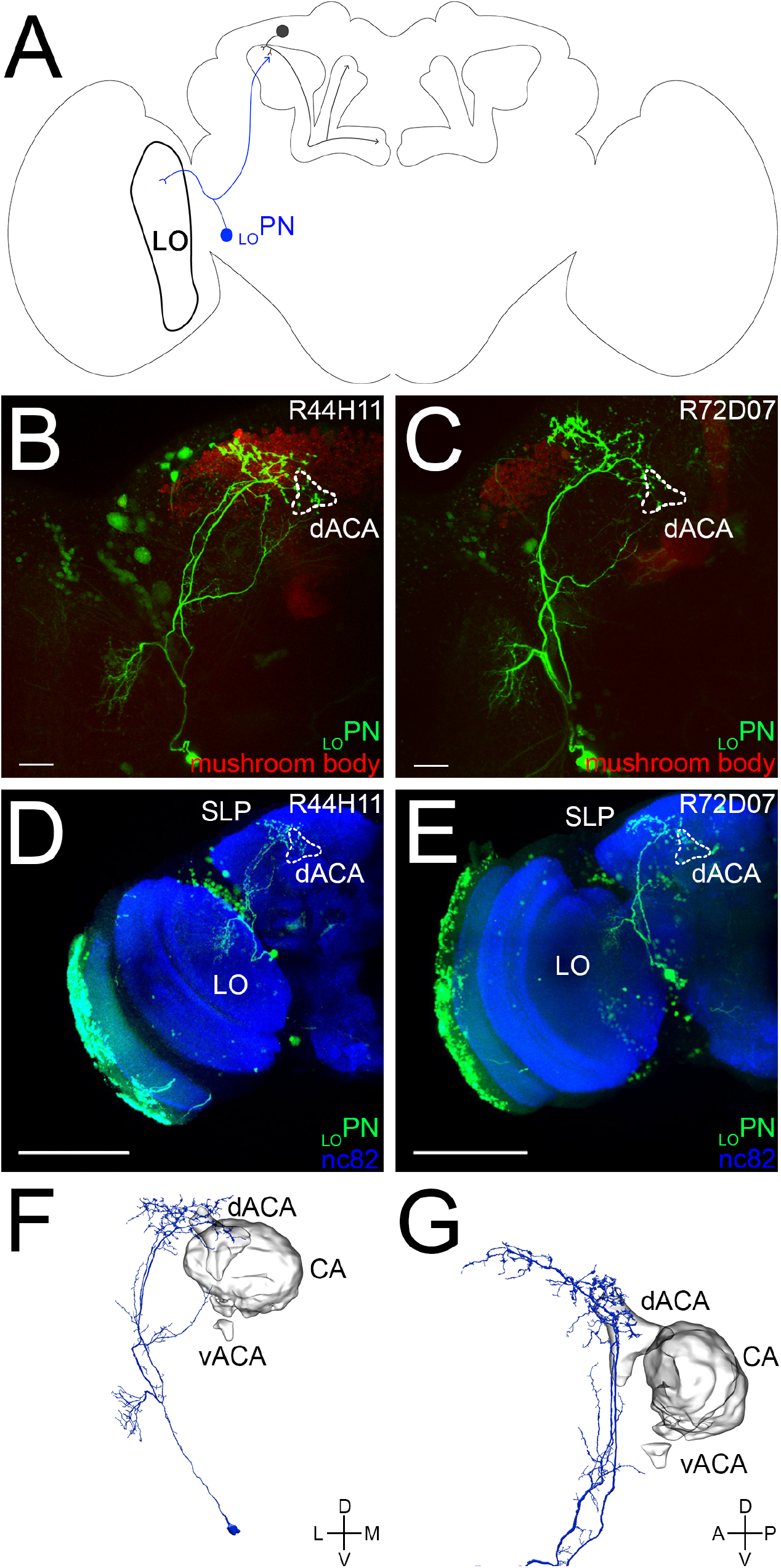
_LO_PN connecting the lobula to the dorsal accessory calyx. (A) A schematic of the *Drosophila* brain shows an α/β_p_ Kenyon cell input neuron — _LO_PN (blue) — projecting from the lobula (LO) to the dorsal accessory calyx. (B-C) _LO_PN (bright green) was identified in the screen using two different transgenic lines (B: *R44H11-GAL4* and C: *R72D07-GAL4);* the neurons photolabeled in each line show an overall similar morphology. (D-E) _LO_PN was photo-labeled using either the *R44H11-GAL4* (D) *R72D07-GAL4* (E) transgenic lines; the samples were fixed, immuno-stained (nc82 antibody, blue) and imaged. The photo-labeled neurons show an overall similar morphology: their somata are located near the optic lobe; they extend dendritic terminals in a small region of the lobula (LO); they extend axonal terminals in the dorsal accessory calyx (dACA) and the superior lateral protocerebrum (SLP). (F-G) A _LO_PN-like neuron was identified in the hemibrain connectome: (F) this neuron projects from the lobula to the dorsal accessory calyx and the superior lateral protocerebrum; (G) its axonal terminals innervate the dorsal accessory calyx (dACA), but not the main calyx (CA) or the ventral accessory calyx (vACA). The following genotypes were used in this figure: (B, D) *yw/yw;MB247-DsRed^unknown^,UAS-C3PA-GFP^unknown^/UAS-C3PA-GFP^attP40^;UAS-C3PA-GFP^attP2^,UAS-C3PA-GFP^VK00005^,UAS-C3PA-GFP^VK00027^/R44H11-GAL4^attP2^*; and (C, E) *yw/yw;MB247-DsRed^unknown^, UAS-C3PA-GFP^attP40^/UAS-C3PA-GFP^unknown^;UAS-C3PA-GFP^attP2^,UAS-C3PA-GFP^VK00005^,UAS-C3PA-GFP^VK00027^/R72D07-GAL4-GFP^attP2^*;. Scale bars are 20 μm (B-C) and 100 μm (D,E).

Therefore, we named this type of neuron “_LO_PN”. The soma of _LO_PN is located in cluster PLP, a region medial to the optic lobe (Figure 2E, 5D-E). _LO_PN extends its dendrites to the lobula and projects its axon to the superior lateral protocerebrum and the dorsal accessory calyx, completely avoiding the main calyx and the ventral accessory calyx (Figure 5D-E). It is worth noting that _LO_PN is very similar to the OLCT1 neuron that has been previously described using a different transgenic line, *R95F09-GAL4;* accordingly, we recovered a strong GRASP signal for that transgenic line (Figure 4L, Table 1; (Yagi et al., 2016)).

We then set out to confirm whether _LO_PN does indeed connect to α/β_p_ Kenyon cells, as surmised from the GRASP results. A previous study has demonstrated that a technique, which combines photo-labeling and dye-labeling tools, can be used to identify the complement of input that individual Kenyon cells receive from the antennal lobe, and the frequency at which individual projection neurons connect to Kenyon cells (Caron et al., 2013). We have modified this technique in order to directly measure the connectivity rate between a given projection neuron and Kenyon cells. In this modified version of the technique, we used a combination of transgenic lines to photolabel the projection neuron of interest — using the same protocol that we described in the previous section — and to dye-label a single Kenyon cell (Figure 6A). We then assessed how many of the dendritic claws formed by the dye-labeled Kenyon cell are connected to a given projection neuron. As a proof of principle, we measured the connectivity rate of the DC3 glomerulus projection neurons to the olfactory Kenyon cells that innervate the main calyx. The connectivity rate of the DC3 projection neurons to the olfactory Kenyon cells has been previously approximated to be 5.1%; this means that there is a 5.1% chance that a given olfactory Kenyon cell claw receives input from the DC3 projection neuron (Caron et al., 2013). Using the modified technique, we found that the DC3 projection neurons connect to Kenyon cells at a connectivity rate of 4.3% (*n* = 30), a value well within the range measured previously (data not shown). We measured the connectivity rates of _LO_PN to α/β_p_ Kenyon cells using the two transgenes we identified in our screen. Using the *R44H11-GAL4* driver line, we found that 4.1 % (*n* = 27) of the claws formed by the α/β_p_ Kenyon cells connect to _LO_PN (Figure 6B-C, H and Table 1). Likewise, using the *R72H07-GAL4* driver line, we found that 3.0 % (*n* = 27) of the claws formed by the α/β_p_ Kenyon cells connect to _LO_PN (Figure 6H and Table 1). Altogether, based on the results obtained using both the GRASP technique and the dye/photo-labeling technique, we conclude that _LO_PN is a true input neuron of the α/β_p_ Kenyon cells and that it conveys information from the lobula, a visual processing center, to the dorsal accessory calyx.

**Figure 6.**
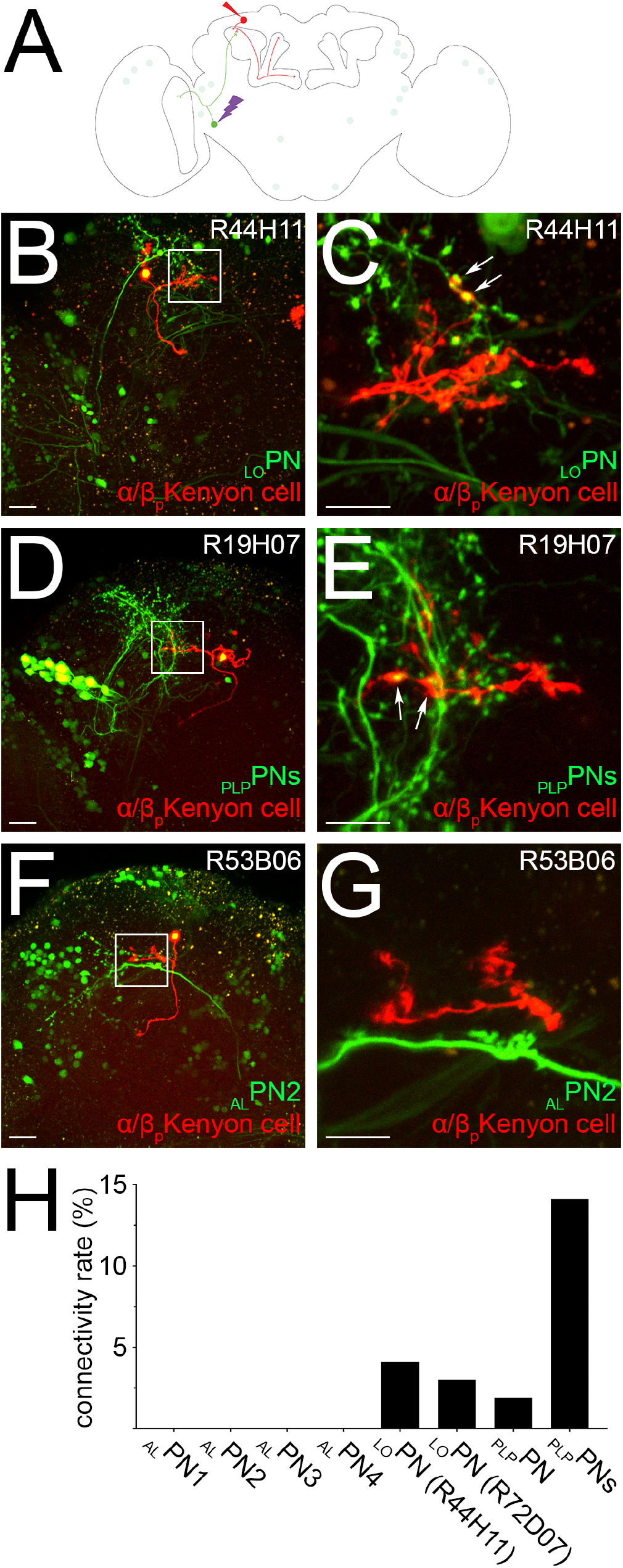
Frequency of connections between the α/β_p_ Kenyon cells and their input neurons. (A) A schematic of the *Drosophila* brain shows how the frequency of connections between the α/β_p_ Kenyon cells (red) and a given input neuron (green), here _LO_PN, was measured. (B-G) A given input neuron was photo-labeled (bright green) and a randomly chosen α/β_p_ Kenyon cell was dye-filled (red). The total number of claws formed by the dye-filled Kenyon cell was counted and claws connecting to the axonal terminals of the photo-labeled neuron were detected (arrows). Such connections were found for _LO_PN (B-C), _PLP_PNs (D-E) but not for the other neurons identified in this study such as _AL_PN2 (F-G). (H) The frequency of connections between α/β_p_ Kenyon cells and a given input neuron was calculated by dividing the number of connections detected (for instance arrows in C and E) by the total number of claws sampled for that particular input neuron. The following genotypes were used in this figure: *yw/yw;UAS-C3PA-GFP^unknown^/UAS-SPA-GFP^attP40^; R13F02-GAL4_AD_^VK00027^,R85D07-GAL4_DBD_^attP2^/R_line* (as indicated in the panel)-*GAL4^attP2^*;. Scale bars are 20 μm (B, D, F) and 10 μm (C, E, G).

### _PLP_PNs connect the posterior lateral protocerebrum to the α/β_p_ Kenyon cells

Photo-labeling of neurons that project to the dorsal accessory calyx using the *R19H07-GAL4* and *R20G07-GAL4* transgenic lines revealed a group of neurons (Figure 7A). In each line, neurons that project from the posterior lateral protocerebrum to the dorsal accessory calyx were photolabeled (Figure 7B-C). Therefore, we named this type of neuron “_PLP_PNs”. The somata of the _PLP_PNs are located in cluster LH, a region near the lateral horn (Figure 7D). We could identify 13 ± 4 _PLP_PNs (*n* = 21) per hemisphere using the *R19H07-GAL4* transgenic line (Figure 7D). The *R20G07-GAL4* transgenic line drives expression at a much lower level, making it difficult to analyze the _PLP_PNs in great detail (Figure 7C). _PLP_PNs extends projections into many brain centers, including the posterior lateral protocerebrum, the superior clamp, the superior lateral protocerebrum and the dorsal accessory calyx, avoiding completely the main calyx and the ventral accessory calyx (Figure 7D). Using the dye/photo-labeling technique described above, we measured the connectivity rate between _PLP_PNs and α/β_p_ Kenyon cells. We found that the _PLP_PNs identified using the *R19H07-GAL4* transgenic line connect at a high rate: we found that 14.1 % (*n* = 24) of the claws formed by the α/β_p_ Kenyon cells connect to the _PLP_PNs (Figure 6D, E, H). We also measured the connectivity rate of individual _PLP_PNs using the same transgenic line and found that, on average, an α/β_p_ Kenyon cell claw has 1.90% (*n* = 32) probability of connecting to a _PLP_PN (Figure 6H and Table 1).

**Figure 7.**
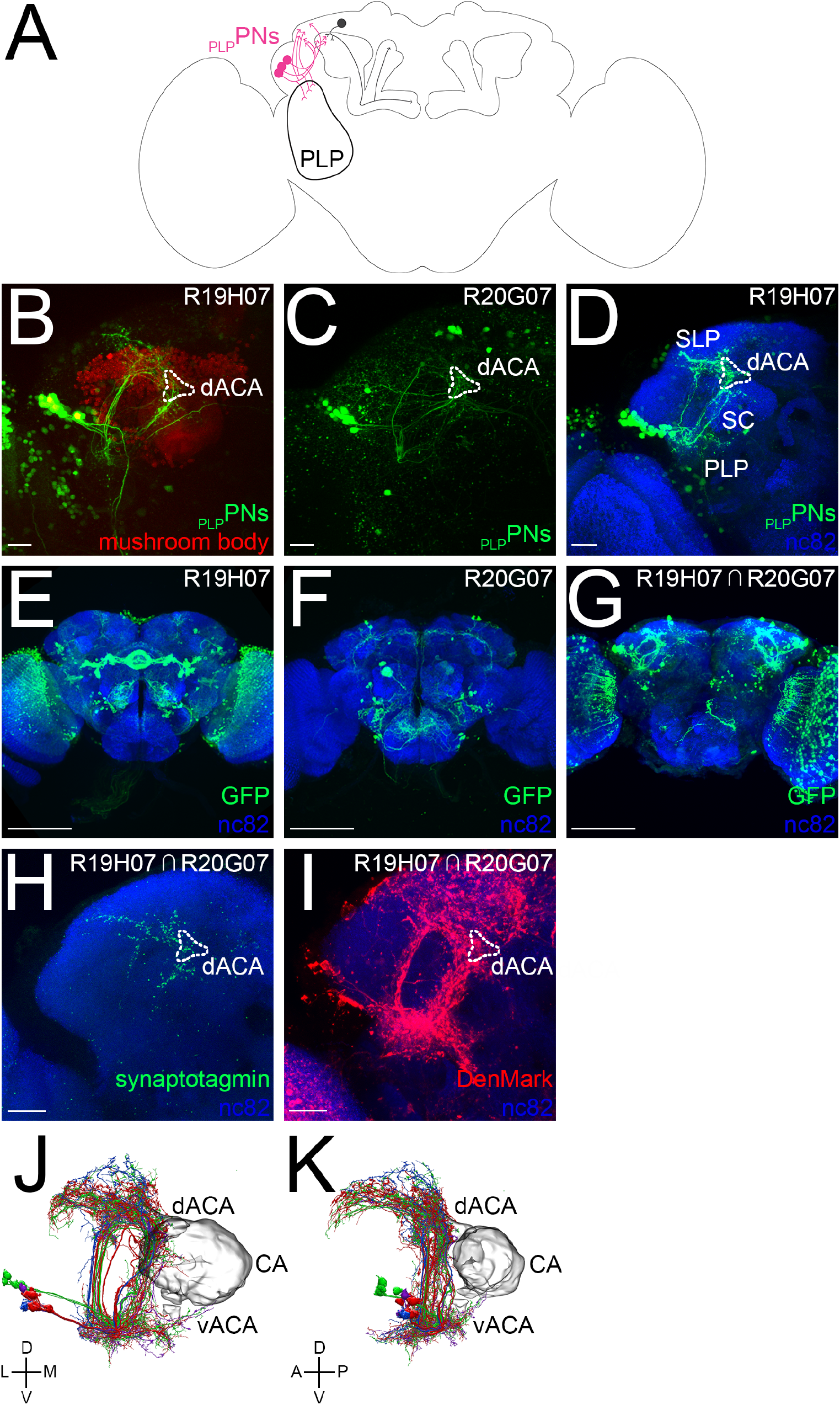
_PLP_PNs connecting the posterior lateral protocerebrum to the dorsal accessory calyx. (A) A schematic of the *Drosophila* brain shows a group of α/β_p_ Kenyon cell input neurons — _PLP_PNs (pink) — projecting from the posterior lateral protocerebrum (PLP) to the dorsal accessory calyx. (B-C) A group of _PLP_PNs (bright green) were identified in the screen using two different transgenic lines (B: *R19H07-GAL4* and C: *R20G07-GAL4).* (D) _PLP_PNs were photolabeled using the *R19H07-GAL4* transgenic line; the sample was fixed, immuno-stained (nc82 antibody, blue) and imaged. The photo-labeled neurons show an overall similar morphology: their somata are located near the lateral horn; they extend projections in the posterior lateral protocerebrum (PLP), the superior clamp (SC), the superior lateral protocerebrum (SLP) and the dorsal accessory calyx (dACA). (E-F) The expression patterns observed for each of the GAL4 lines are broad and include many neurons; (E) the *R19H07-GAL4* line drives expression strongly in many neurons, including the _PLP_PNs, (F) whereas the *R20G07-GAL4* line drives expression weakly in fewer neurons, including the _PLP_PNs. These images were obtained from the FlyLight website. (G) A split-GAL4 transgenic line driving expression strongly in the _PLP_PNs was engineered using the *R19H07* promoter region to drive the GAL4_AD_ domain and the *R20G07* promoter region to drive the GAL4_DBD_. (H-I) The split-GAL4 line was used to drive the expression of the presynaptic marker synaptotagmin-fused GFP (H) or the post-synaptic maker DenMark (I). (J-K) _PLP_PN-like neurons were identified in the hemibrain connectome: (J) these neurons project from the posterior lateral protocerebrum to the dorsal accessory calyx, the superior lateral protocerebrum and the superior clamp; (K) their axonal terminals innervate the dorsal accessory calyx (dACA), but not the main calyx (CA) or the ventral accessory calyx (vACA). The following genotypes were used in this figure: (B, D) *yw/yw; MB247-DsRed^unknown^, UAS-C3PA-GFP^unknown^/UAS-C3PA-GFP^attP40^;UAS-C3PA-GFP^attP2^, UAS-C3PA-GFP^VK00005^, UAS-C3PA-GFP^VK00027^/R19H07-GAL4^attP2^*; (C) *yw/yw; UAS-C3PA-GFP^unknown^/CyO;UAS-C3PA-GFP^attP2^, R20G07-GAL4^attP2^/MKRS;* (G) *yw/yw; MB247-DsRed^unknown^, UAS-C3PA-GFP^unknown^/CyO; UAS-C3PA-GFP^attP2^, UAS-C3PA-GFP^VK00005^,UAS-C3PA-GFP^VK00027^/R20G07-GAL4_DBD_^attP2^,R19H07-GAL4_AD_^VK00027^*; (H) *yw/yw;Sp/CyO; R20G07-GAL4_DBD_^attP2^,R19H07-GAL4_AD_^VK00027^/UAS-synaptotagmin::GFP^unknown^*; (I) *yw/yw,UAS-DenMark^2^/CyO; R20G07-GAL4_DBD_^attP2^,R19H07-GAL4_AD_^VK00027^/Tm2*;. Scale bars are 20 μm (A-D), 100 μm (E-G) and 20 μm (H-I).

To determine whether the _PLP_PNs identified in the *R19H07-GAL4* and *R20G07-GAL4* transgenic lines are the same or different group of neurons, we used the split-GAL4 technique (Luan et al., 2006). In this technique, the GAL4 transcription factor is split into two complementary fragments — the GAL4_AD_ and GAL4_DBD_ domains — that are both transcriptionally inactive when expressed alone; when both fragments are expressed in the same cell, GAL4 can be reconstituted and recovers its transcriptional activity. _PLP_PNs were visible in flies that carry the *R20G07-GAL4_DBD_* and *R19H07-GAL4_AD_* transgenes, thus confirming that both lines are expressed in the same group of _PLP_PNs (Figure 7E-G). This result revealed, even more clearly, that individual _PLP_PNs project in many different brain centers (Figure 7G). In order to distinguish the axonal terminals from the dendritic arbors, we used the split-GAL4 combination of transgenic lines (the *R20G07-GAL4_DBD_* and *R19H07-GAL4_AD_* transgenic lines) to drive the expression of the presynaptic marker synaptotagmin in the _PLP_PNs (Figure 7H). We found that the projections extending into the superior lateral protocerebrum, superior clamp and dorsal accessory calyx contain presynaptic terminals. Similarly, when the expression of the postsynaptic marker DenMark was driven specifically in the _PLP_PNs, we found that all of the projections made by the _PLP_PNs contain postsynaptic terminals (Figure 7I). However, the projections extending into the posterior lateral protocerebrum are the only projections formed by _PLP_PNs that contain only post-synaptic terminals and no pre-synaptic terminals (Figure 7H-I).

To identify the neurons that project to the post-synaptic terminals formed by the _PLP_PNs in the posterior lateral protocerebrum, a poorly characterized visual processing center, we used the targeted photo-labeling technique described above (Figure 8A). In short, we used a combination of transgenes that, in concert, drive the expression of PA-GFP in all neurons (the *N-Synaptobrevin-QF* and *QUAS-PA-GFP* transgenes) and the expression of tdTomato in the _PLP_PNs (using the *R19H07-GAL4_AD_, R20G07-GAL4_DBD_* and *UAS-tdTomato* transgenes). Guided by the expression of tdTomato, we targeted the post-synaptic terminals formed by _PLP_PNs in the posterior lateral protocerebrum with high-energy light (Figure 8B, C). Upon photo-labeling, two types of neuron were clearly photo-labeled: the _PLP_PNs — showing that the photo-activation was specific to these neurons — and neurons projecting from the ventral medulla (Figure 8D). These photolabeled neurons project into deeper layers of the medulla (Figure 8E). Altogether, from this set of experiments, we conclude that _PLP_PNs are one of the major input neurons of the α/β_p_ Kenyon cells and that they convey information from the posterior lateral protocerebrum, and possibly from the ventral medulla, to the dorsal accessory calyx.

**Figure 8.**
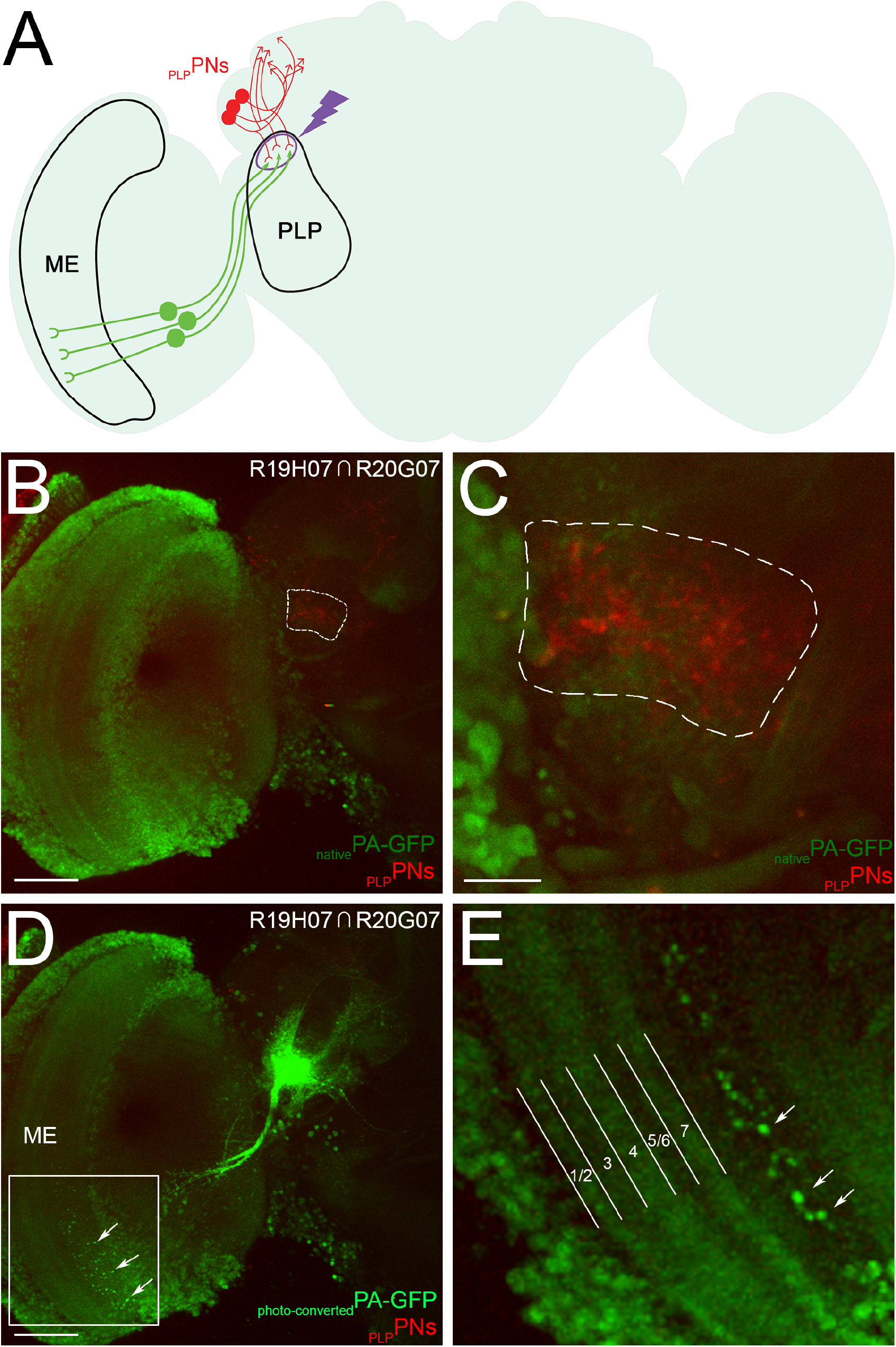
Identification of the _PLP_PNs input neurons using photo-labeling. (A) A schematic of the *Drosophila* brain shows how neurons projecting from the ventral medulla (ME) were identified by photo-labeling the dendritic terminals formed by the _PLP_PNs in the posterior lateral protocerebrum (PLP). (B-C) All neurons express PA-GFP (light green) and the _PLP_PNs express tdTomato (red) (B). The dendritic terminals formed by the _PLP_PNs in the posterior lateral protocerebrum are visible in the red channel, and the region outlined by the white dashed line was targeted for photo-labeling (C). (D-E) Upon photo-activation of PA-GFP in the posterior lateral protocerebrum, neurons projecting from the ventral lobula were identified (D); these neurons extend dendritic arbors (white arrows) in a layer of the lobula deeper than the well-characterized layers 1-7 (E). The following genotype is used in this figure: *yw/yw; QUAS-C3PA-GFP^unknown^, QUAS-SPA-GFP^unknown^/UAS-tdtomato^attP40^; R20G07-GAL4_DBD_^attP2^,R19H07-GAL4_AD_^VK00027^/N-Synaptobrevin-QF;.* Scale bars are 50 μm (B, D) and 10 μm (C).

### _AL_PNs do not contribute major input to the α/β_p_ Kenyon cells

Photo-labeling using the transgenic lines that displayed weak GRASP signal — these are the *R30E11-GAL4, R31C03-GAL4* and *R53B06-GAL4* lines — or no GRASP signal — these are the *R11F07-GAL4, R11F08-GAL4* and *R12C04-GAL4* lines — led to the identification of a third type of neuron (Figure 9). The somata of these neurons are all located in AL cluster, a region near the antennal lobe. Although each of these neurons has a distinct overall morphology, they all project from the antennal lobe to the superior lateral protocerebrum and extend their axons in a region near the dorsal accessory calyx. Therefore, we named this type of neuron “_AL_PNs”.

**Figure 9.**
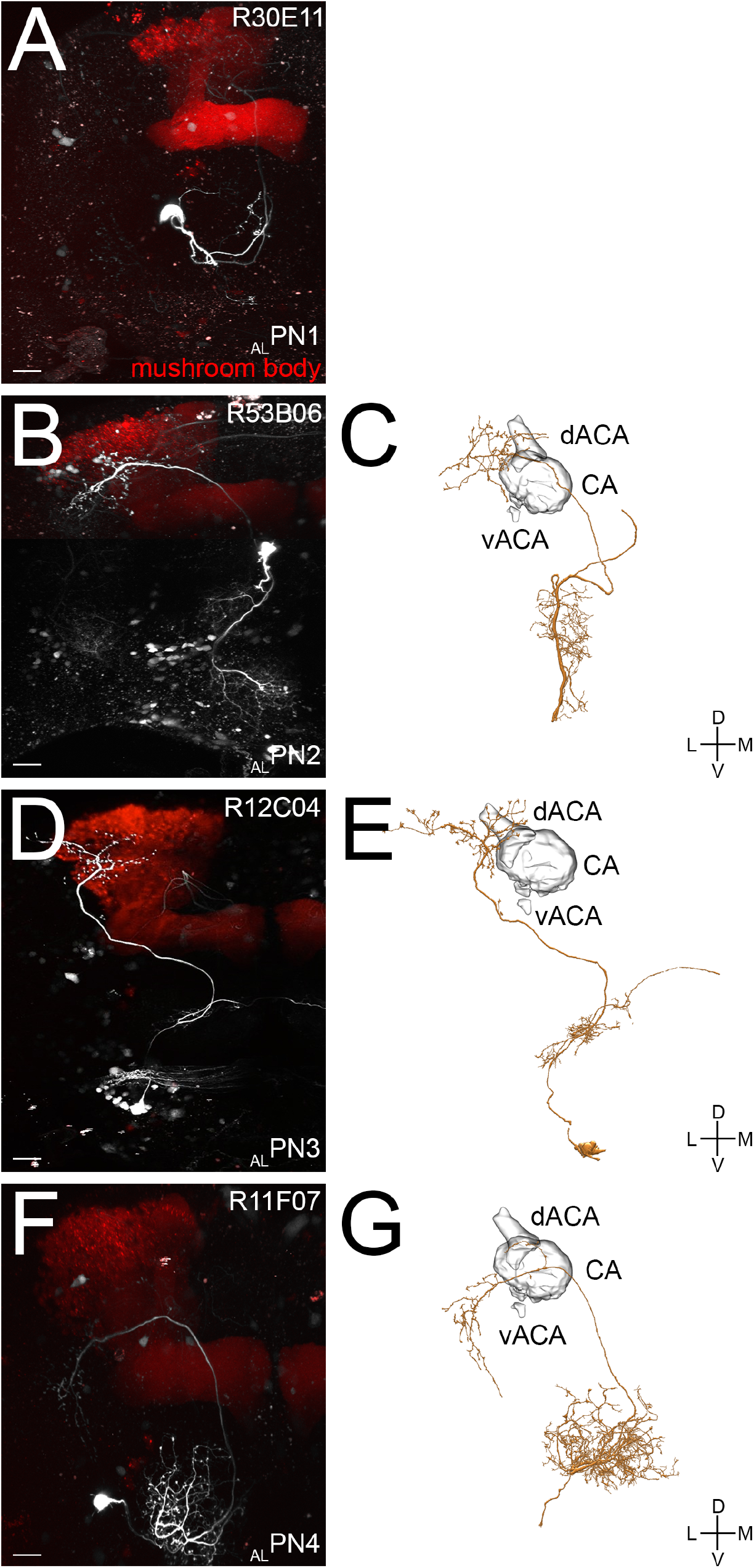
_AL_PN projecting from the antennal lobe to region near the dorsal accessory calyx. (A, B, D, F) Four different neurons projecting from the antennal lobe to a region near the dorsal accessory calyx were identified using different transgenic lines (A: *R30E11-GAL4,* B: *R53B06-GAL4,* C: *R12C04-GAL4* and D: *R11F071-GAL4);* all the photo-labeled neurons project from the antennal lobe but each neuron has a distinct morphology. (C, E, G) Three of the four ALPN-like neurons were identified in the hemibrain connectome. (A) _AL_PN1 extends its dendrites in the posterior antennal lobe and projects its axons in the superior lateral protocerebrum. (B-C) _AL_PN2 extends its dendrites into the column region of the posterior antennal lobe, a region known to be activated by high humidity, as well as in the sub-esophageal ganglion, a gustatory processing center, and projects its axons in the superior lateral protocerebrum. (D-E) _AL_PN3 extends its dendrites into the arm region of the posterior antennal lobe, a region known to be activated by low humidity and projects its axons in the superior lateral protocerebrum. (F-G) _AL_PN4 extends its dendrites broadly in the anterior antennal lobe, an olfactory processing center, and projects its axons in the posterior lateral protocerebrum. The following genotypes were used in this figure: *yw/yw;MB247-DsRed^unknown^,UAS-C3PA-GFP^unknown^/UAS-C3PA-GFP^attP40^;UAS-C3PA-GFP^attP2^,UAS-C3PA-GFP^VK00005^,UAS-C3PA-GFP^VK00027^/R_line-GAL4^attP2^(as* indicated on the panel);. Scale bars are 20 μm.

The neurons photo-labeled using the *R30E11-GAL4* and *R31C03-GAL4* driver lines show a nearly identical morphology: in each line, a single neuron that extends its dendrites into the posterior antennal lobe and project its axons into the superior lateral protocerebrum was visible (Figure 9A). We named this neuron “_AL_PN1”. It is worth noting that _AL_PN1 is very similar to the thermosensitive AC neuron that has been previously described (Shih and Chiang, 2011). Using a combination of photo-labeling and dye-tracing techniques, as described above, we measured the connectivity rate between _AL_PN1 and α/β_p_ Kenyon cells. We could not detect any connections between _AL_PN1 and α/β_p_ Kenyon cells (*n* = 26), suggesting that _AL_PN1 is not a dorsal accessory calyx input neuron (Figure 6H, Table 1). Similarly, the neuron photo-labeled using the *R53B06-GAL4* line extends its dendrites into the column region of the posterior antennal lobe, a region known to be activated by high humidity, and the sub-esophageal ganglion, a gustatory processing center (Figure 9B). We named this neuron “_AL_PN2”. Again, we could not detect any connections between _AL_PN2 and α/β_p_ Kenyon cells (*n* = 30), suggesting that _AL_PN2 is not a dorsal accessory calyx input neuron (Figure 6F-H, Table 1). Thus, _AL_PN1 and _AL_PN2 are most likely not contributing major input to the α/β_p_ Kenyon cells.

We extended our analysis to the transgenic lines that displayed no GRASP signal. The neurons photo-labeled using the *R11F08-GAL4* and *R12C04-GAL4* driver lines show an overall similar morphology: in each line, a single neuron that extends its dendrites into the arm region of the posterior antennal lobe, a region known to be activated by low humidity, was visible (Figure 9D). We named this neuron “_AL_PN3”. Not surprisingly, we could not detect any connections between _AL_PN3 and α/β_p_ Kenyon cells (*n* = 26) suggesting that _AL_PN3 does not provide input into the dorsal accessory calyx (Table 1). Similarly, the neuron photo-labeled using the *R11F07-GAL4* line extends its dendrites broadly throughout the anterior antennal lobe, an olfactory processing center (Figure 9F). We named this neuron “_AL_PN4”. Again, we could not detect any connections between _AL_PN4 and α/β_p_ Kenyon cells (*n* = 25), suggesting that _AL_PN4 does not provide input into the dorsal accessory calyx (Table 1). Thus, we could confirm that the transgenic lines that displayed no GRASP signal do not appear to provide major input to the α/β_p_ Kenyon cells. Altogether, these results show that the third type of neuron identified in our screen — _AL_PNs — project to a region close to the dorsal accessory calyx but are most likely not presynaptic to the α/β_p_ Kenyon cells.

### _LO_PN and _PLP_PNs are major input neurons of the α/β_p_ Kenyon cells

Our results suggest that the _LO_PN and _PLP_PNs that we identified in our screen connect to the α/β_p_ Kenyon cells and that, together, these two types of projection neuron represent a large fraction of the input neurons of the dorsal accessory calyx. Our results also suggest that the _AL_PNs do not connect to the α/β_p_ Kenyon cells, as another study suggested (Yagi et al., 2016). Thus, we conclude that the dorsal accessory calyx is anatomically poised to receive information primarily from the lobula and the posterior lateral protocerebrum, two visual processing centers. We verified whether these findings corroborate with the recently released *Drosophila* hemibrain connectome (Xu et al., 2020). Using the Neuprint 1.0.1 platform, we focused our attention on the 60 reconstructed α/β_p_ Kenyon cells that have been fully traced. Altogether, these α/β_p_ Kenyon cells receive input from a large number of neurons but most of these synapses are most likely axoaxonic synapses as they are located along the α and β lobes. These input neurons are primarily other Kenyon cells, various dopaminergic neurons as well as neurons known to connect broadly to all Kenyon cells such as the APL neuron and the DPM neuron (data not shown). We focused our attention on the 141 input neurons that connect to the α/β_p_ Kenyon cells in the dorsal accessory calyx. Not surprisingly, in accordance with the results obtained from our *en masse* photo-labeling experiment, the cell bodies of these input neurons can be divided into seven different clusters and the ratio of neurons belonging to a given cluster is largely consistent between both analyses (Figure 10A).

**Figure 10.**
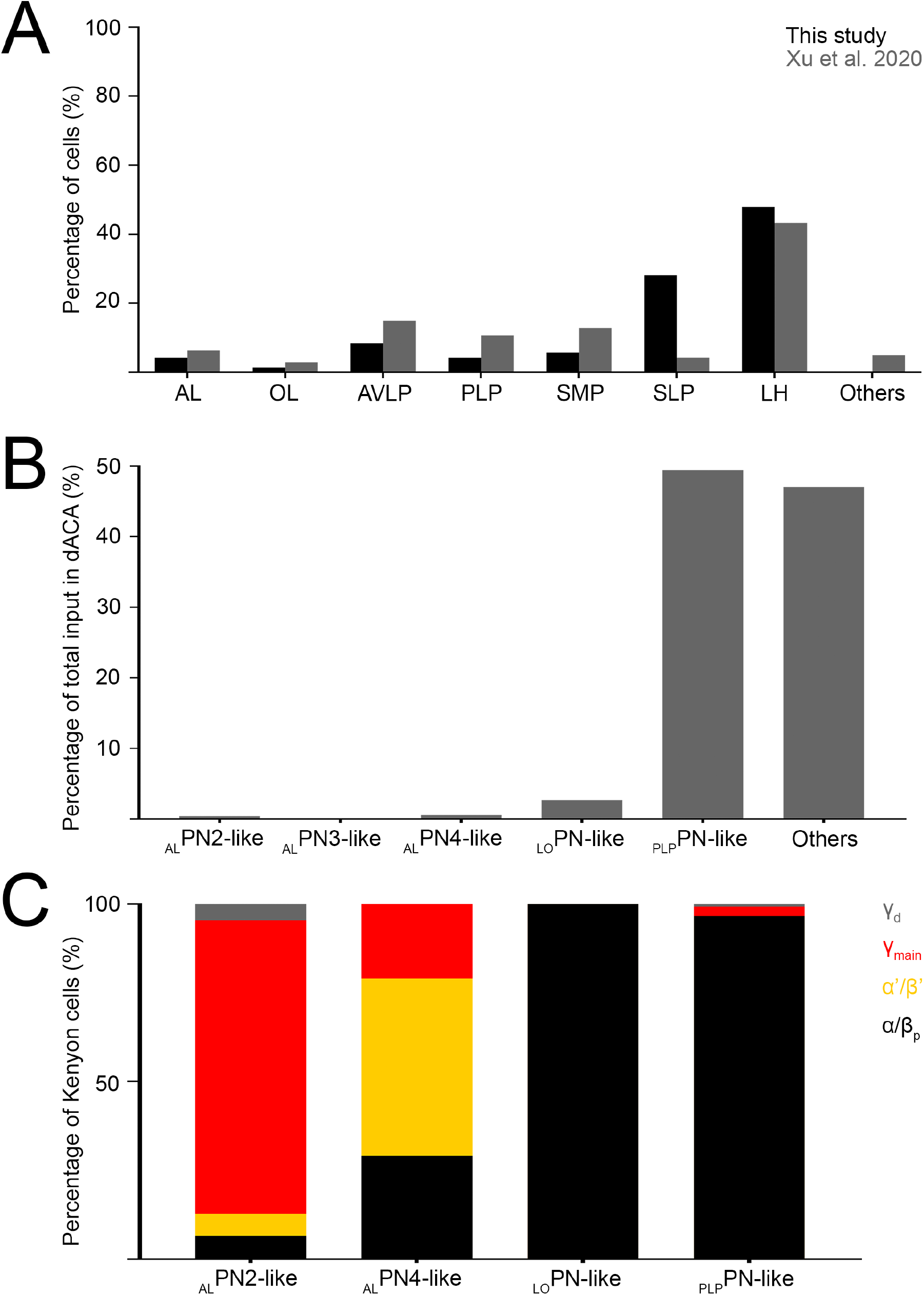
Identification of the projection neurons connecting to the α/β_p_ Kenyon cells in the dorsal accessory calyx in the connectome. (A-C) The 60 α/β_p_ Kenyon cells reconstructed in the *Drosophila* hemibrain connectome receive input from 141 neurons in the dorsal accessory calyx. (A) These input neurons can be divided into seven clusters based on the location of their cell bodies: the antennal lobe (AL), optic lobe (OL), anterior ventral lateral protocerebrum (AVLP), lateral horn (LH), superior lateral protocerebrum (SLP), superior medial protocerebrum (SMP) and posterior lateral protocerebrum (PLP) clusters; neurons with somata located outside these clusters are defined as “others”. The percentage of neurons found in each cluster (grey) is similar to the one measured using the *en masse* photo-labeling technique reported in this study (black). (B) Among these 141 input neurons, neurons morphologically similar to _AL_PNs, _LO_PN and _PLP_PNs were identified; the percentage of input α/β_p_ Kenyon cells receive in the dorsal accessory calyx from these different types of neuron varies across types. (C) ALPN-like, _LO_PN-like and _PLP_PN-like neurons connect to different types of Kenyon cell; whereas _AL_PNs connect mostly to olfactory Kenyon cells (namely γ_main_ (red) and α’/β’ (yellow) Kenyon cells), _LO_PN-like and _PLP_PN-like neurons connect mostly to α/β_p_ Kenyon cells (black). Very few connections to γ_d_ Kenyon cells (grey) are found.

Interestingly, the LH cluster is the most numerous cluster: our study identified 34 ± 12 neurons (*n* = 22) in this cluster — including the 13 ± 4 _PLP_PNs (*n* = 21) — whereas the connectome shows a total of 61 input neurons with cell bodies located in the lateral horn (Figure 10A). Among these neurons, we identified 13 that are morphologically highly similar to _PLP_PNs (Figure 7J-K, Table 2). We found that these _PLP_PN-like neurons connect to 59 of the 60 reconstructed α/β_p_ Kenyon cells, representing 49.40% of the input all α/β_p_ Kenyon cells receive in the dorsal accessory calyx (Figure 10B and Table 2). _PLP_PNs-like neurons connect mostly to α/β_p_ Kenyon cells but also form a few connections to γ_d_ and γ_main_ Kenyon cells, reinforcing the observation that the dorsal accessory calyx input neurons and the ventral accessory calyx input neurons form two parallel pathways (Figure 10C). In our study, we identified neurons connecting the ventral medulla to the _PLP_PNs (Figure 8) but, because the hemibrain connectome does not include the medulla, we could not identify fully traced neurons similar to these _PLP_PNs input neurons. Additionally, our study identified 1 ± 2 neurons (*n* = 22) in the OL cluster — including _LO_PN — whereas the connectome shows a total of 4 input neurons with cell bodies located in the optic lobe (Figure 10A). Among these neurons, we identified a neuron very similar to _LO_PN (Figure 5F-G). We found that this _LO_PN-like neuron connects to 11 of the 60 reconstructed α/β_p_ Kenyon cells, representing 2.66% of the input all α/β_p_ Kenyon cells receive in the dorsal accessory calyx (Figure 10B and Table 2). As we found in our study, this _LO_PN-like neuron does not connect to any other types of Kenyon cell, forming a pathway parallel to the one conveying visual information to the ventral accessory calyx.

**Table 2.** Input neurons of the dorsal accessory calyx as identified in the *Drosophila melanogaster* hemibrain connectome. The neurons connecting to α/β_p_ Kenyon cells in the dorsal accessory calyx — and their corresponding neurons, as described in this study — are listed; the numbers in parentheses indicate the number of neurons reported for each type. The identity number can be used to visualize each of these neurons using the Neuprint platform. Also listed are the number of α/β_p_ Kenyon cells connected to a given input neuron, as reported in Neuprint v1.0.1, as well as the percentage of input these neurons provide to the α/β_p_ Kenyon cells in parantheses.

| Type of neuron<br>(this study) | Type of neuron<br>(Xu et al., 2020) | Neuprint ID number | Number of $\alpha/\beta_p$ Kenyon<br>cells connected |
| --- | --- | --- | --- |
| $LOPN$ (1) | PVL02b_pct (1) | 419908058 | 11 (2.66%) |
| $PLPN$ (13) | PDL20y_pct (5) | 479969106 | 23 (6.72%) |
|  |  | 5813019543 | 10 (2.60%) |
|  |  | 450597298 | 7 (2.75%) |
|  |  | 389536926 | 13 (6.06%) |
|  |  | 5813009790 | 2 (0.39%) |
|  | PDL23g_pct (4) | 5813011738 | 20 (8.75%) |
|  |  | 480310599 | 4 (0.27%) |
|  |  | 454335501 | 15 (5.01%) |
|  |  | 295461570 | 2 (0.57%) |
|  | PDL20ob_pct (2) | 5813011717 | 9 (3.07%) |
|  |  | 511634742 | 5 (0.33%) |
|  | PDL20z_pct (2) | 512023157 | 25 (8.78%) |
|  |  | 480997558 | 15 (4.12%) |
| $ALPN2$ (1) | VP multi adPN mALT | 5813063239 | 2 (0.39%) |
| $ALPN4$ (1) | olfactory multi lvPN mALT | 1037293275 | 10 (0.54%) |
| N/A (1) | PVL10a_pct | 479935033 | 50 (22.93%) |

Finally, both studies identified a number of neurons with cell bodies located near the antennal lobe: using *en masse* photo-labeling, we identified a total of 3 ± 3 neurons (*n* = 22) in the AL cluster, whereas nine such neurons are found in the connectome. Among these neurons, we recognized three of the four _AL_PNs we characterized: an _AL_PN2-like neuron and an _AL_PN4-like neuron that represents, respectively, 0.39% and 0.54% of the input that α/β_p_ Kenyon cells receive in the dorsal accessory calyx (Figure 9C, G and Table 2). We could not recognize an ALPN1-like neuron in the connectome but we identified an ALPN3-like neuron (Figure 9E, Table 2). The _AL_PN3-like neuron does not connect to any Kenyon cells. Together, the nine antennal lobe input neurons — including the _AL_PN2-like and _AL_PN4-like neurons — represent less than 2% of the input that α/β_p_ Kenyon cells receive in the dorsal accessory calyx (Figure 10B). Interestingly, most of the remaining input α/β_p_ Kenyon cells receive in the dorsal accessory calyx (22.93% of the total input α/β_p_ Kenyon cells receive in the dorsal accessory calyx) is from a single neuron projecting from many brain regions including the main calyx and the pedunculus, two regions of the mushroom body, as well as the superior lateral protocerebrum (Table 2). Altogether, these observations confirm that _LO_PN-like and _PLP_PN-like neurons are the major input neurons that connect to the α/β_p_ Kenyon cells in the dorsal accessory calyx. Thus, the dorsal accessory calyx receives most of its input from visual processing centers — the posterior lateral protocerebrum and the lobula — and is thus anatomically poised to process mostly, if not strictly, visual information.

## Discussion

In this study, we identified and characterized neurons projecting to the dorsal accessory calyx of the mushroom body and show that these neurons are presynaptic to the α/β_p_ Kenyon cells (Figure 11). Using a combination of genetic and anatomical techniques, we could distinguish two different types of projection neuron: _LO_PN projecting from the lobula — an area of the optic lobe processing visual features such as shape and motion — and the _PLP_PNs projecting from the posterior lateral protocerebrum. Although the posterior lateral protocerebrum remains poorly characterized in *D. melanogaster,* evidence from other insects shows that this brain region receives input from the optic lobe (Paulk et al., 2008, 2009). Interestingly, we found that the dendrites formed by the _PLP_PNs in the posterior lateral protocerebrum are in close proximity to neurons that project from the ventral medulla. Based on our results — and considering insights from the connectome — we estimate that _LO_PNs and _PLP_PNs account for half of total input that α/β_p_ Kenyon cells receive in the dorsal accessory calyx. _LO_PNs and _PLP_PNs do not extend axonal terminals into the ventral accessory calyx, the other calyx known to receive visual input, but rather extend axonal terminals into the dorsal accessory calyx and into the superior lateral protocerebrum. Likewise, the α/β_p_ Kenyon cells do not connect to the visual projection neurons that are associated with the ventral accessory calyx (Vogt et al., 2016). These findings suggest that the visual system is connected to the mushroom body via two parallel pathways: the α/β_p_ Kenyon cells receive input from the lobula and the posterior lateral protocerebrum, whereas the γ_d_ Kenyon cells receive input directly from the medulla. Further functional studies are necessary to determine what kind of visual information is processed by the α/β_p_ Kenyon cells.

**Figure 11.**
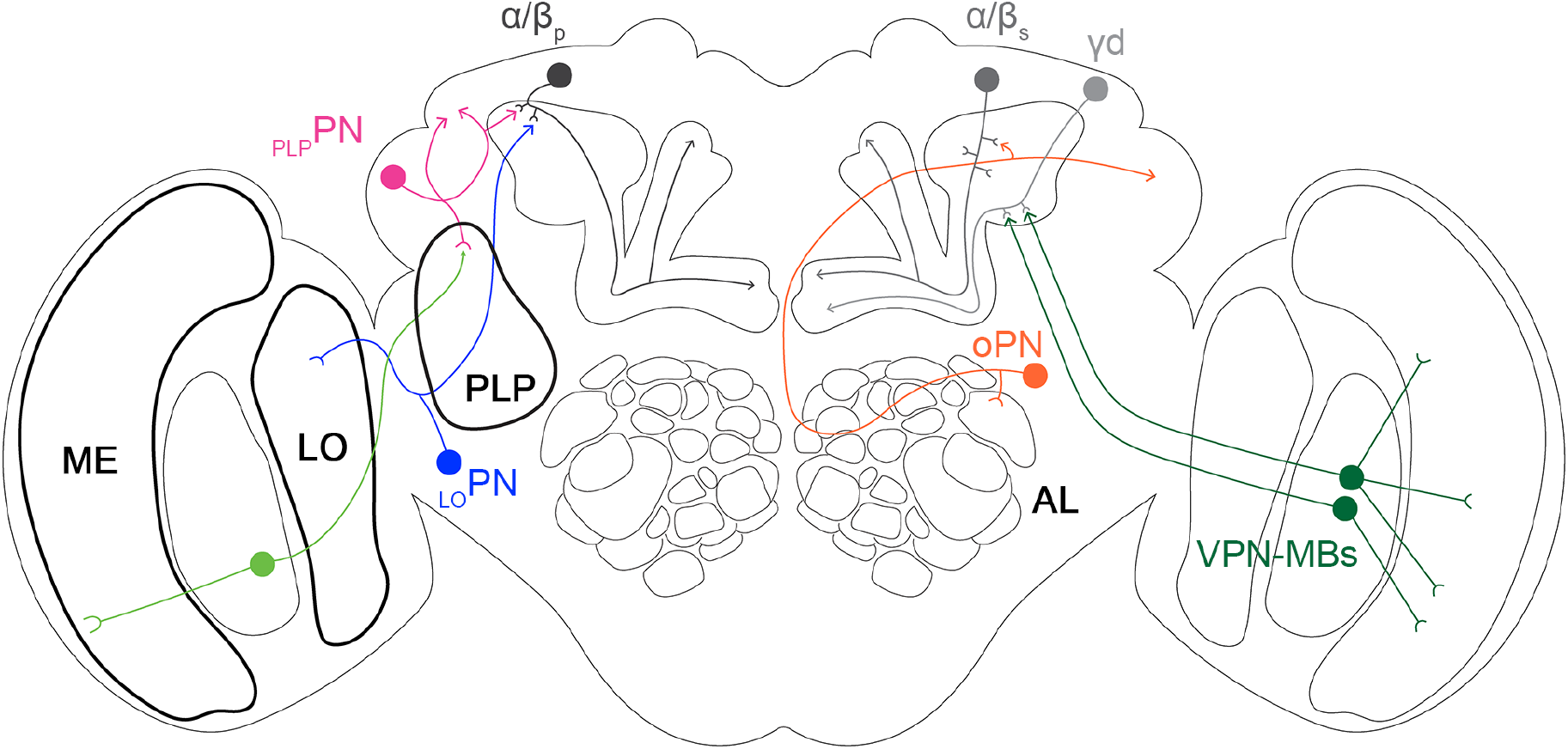
Visual input into the mushroom body through two parallel pathways. The main calyx (CA) of the mushroom body receives input from the olfactory system: neurons projecting from the antennal lobe (orange) form synapses with the Kenyon cells associated with the main calyx, such as the α/β_s_ Kenyon cells. The dorsal accessory calyx (dACA) and the ventral accessory calyx (vACA) receive input from the visual system: a neuron projecting from the lobula (_LO_PN, blue) forms synapses with the α/β_p_ Kenyon cells associated with the dorsal accessory calyx, whereas neurons projecting from the medulla (_ME_PNs, green) form synapses with the γ_d_ Kenyon cells associated with the ventral accessory calyx. The dorsal accessory calyx also receives input from the posterior lateral protocerebrum via the _PLP_PNs (pink), which are connected to neurons projecting from the ventral medulla (green).

In *Drosophila melanogaster*, the mushroom body has long been studied as an olfactory processing center. However, evidence from many insects, including the honeybee *Apis mellifera,* shows that the mushroom body integrates sensory information across different modalities. In honeybees, the input region of the mushroom body, also called the calyx, is divided into different layers and each layer receives input from either the olfactory or visual system (Gronenberg, 2001). Because the dendrites of Kenyon cells are also restricted to specific layers, it has been suggested that, in the honeybee, multisensory integration does not occur at the level of individual Kenyon cells, but rather at the population level (Ehmer and Gronenberg, 2002). Although the honeybee mushroom body differs greatly from the *Drosophila* mushroom body — it contains about a hundred times as many Kenyon cells and its input region is divided in multiple complex layers — it appears that both mushroom bodies share a common fundamental connectivity principle: the segregation of input based on sensory modality. This connectivity mechanism is immediately apparent in the structural organization of the *Drosophila melanogaster* mushroom body: the Kenyon cells receiving input from the olfactory system all extend their dendrites into the main calyx, whereas as the Kenyon cells receiving input from the visual system extend their dendrites either in the dorsal accessory calyx or the ventral accessory calyx. Many studies have demonstrated that the Kenyon cells that process olfactory information — those associated with the main calyx — integrate input broadly across the different types of olfactory projection neuron (Caron et al., 2013; Zheng et al., 2018). Interestingly, it appears that the Kenyon cells that process visual information are wired differently.

*We* have a thorough understanding of how olfactory Kenyon cells integrate input from the antennal lobe: most Kenyon cells receive, on average, input from seven projection neurons and the projection neurons connecting to the same Kenyon cell share no apparent common features (Caron et al., 2013; Zheng et al., 2018). Theoretical studies have shown that this random-like connectivity pattern enables the mushroom body to form sparse and decorrelated odor representations and thus maximizes learning (Litwin-Kumar et al., 2017). Randomization of sensory input is a connectivity pattern that is well suited for representing olfactory information — as an odor is encoded based on the ensemble of olfactory receptors it activates — and might not be suitable for representing visual information. Indeed, our results suggest that that specific visual features — the signals processed by the medulla and the ones processed by the lobula and the posterior lateral protocerebrum — need to be represented by two separate subpopulations of Kenyon cells. This observation mirrors anatomical studies of the honeybee brain: the neurons projecting from the lobula terminate in a different layer than the neurons projecting from the medulla (Ehmer and Gronenberg, 2002). This arrangement might be essential to preserve distinct visual features when forming associative memories. Functional and behavioral studies are required to determine whether indeed the mushroom body represents multisensory stimuli in this manner.

## MATERIALS AND METHODS

### Fly stocks

Flies *(Drosophila melanogaster)* were raised on standard cornmeal agar medium and maintained in an incubator set at 25°C, 60% humidity with a 12 hours light / 12 hours dark cycle (Percival Scientific, Inc.). Crosses were set up and reared under the same conditions, but the standard cornmeal agar medium was supplemented with dry yeast. The strains used in this study are described in the table below.

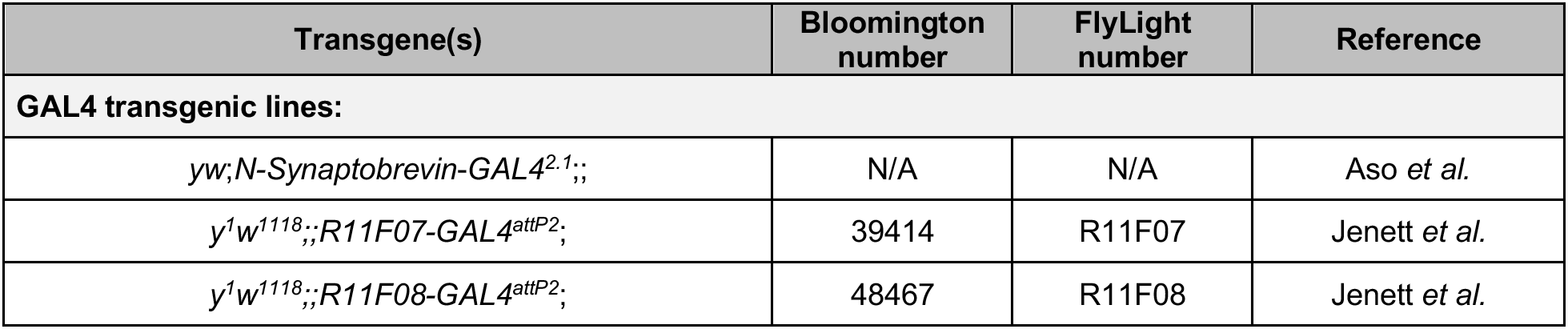
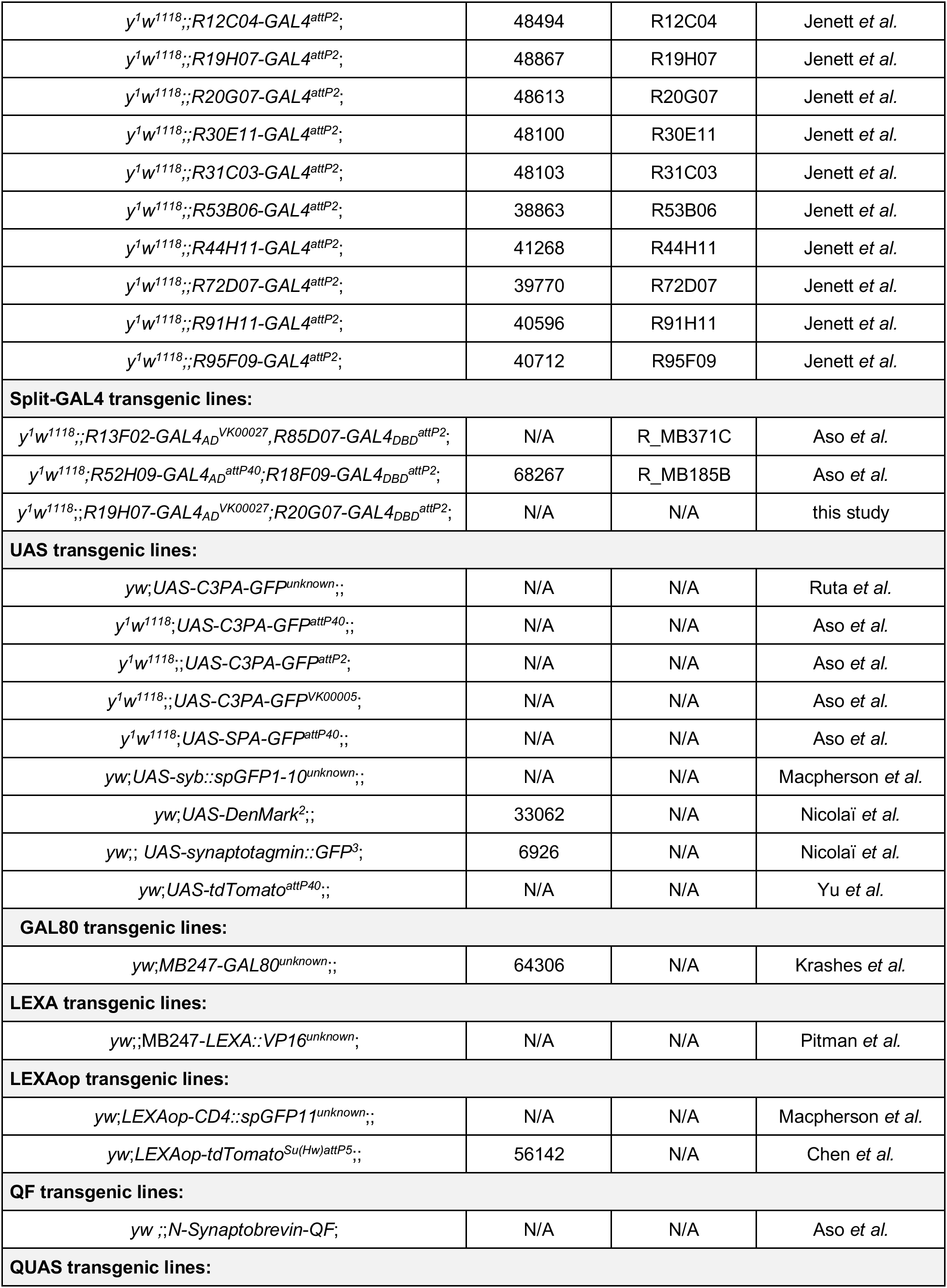
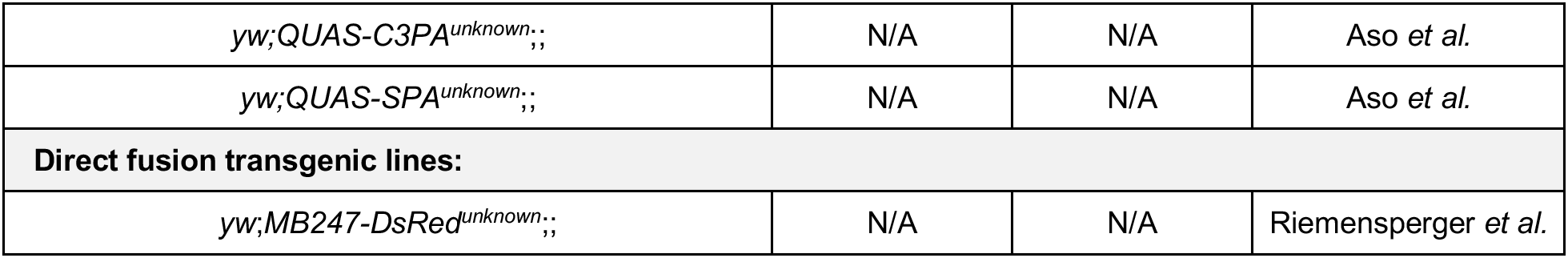

### Photo-labeling neurons using PA-GFP

Two to six-day-old flies were used for all photo-labeling experiments. The protocol was largely based on the one developed in a previous study (Aso et al., 2014). In short, brains were dissected in saline (108 mM NaCl, 5 mM KCl, 5 mM HEPES, 5 mM Trehalose, 10 mM Sucrose, 1 mM NaH_2_PO_4_, 4 mM NaHCO_3_, 2 mM CaCl_2_, 4 mM MgCl_2_, pH≈7.3), treated for 1 minute with 2 mg/ml collagenase (Sigma-Aldrich) and mounted on a piece of Sylgard placed at the bottom of a Petri dish. Photo-labeling and image acquisition were performed using an Ultima two-photon laser scanning microscope (Bruker) with an ultrafast Chameleon Ti:Saphirre laser (Coherent) modulated by Pockels Cells (Conotopics). For photo-labeling, the laser was tuned to 710 nm with an intensity of 5-30 mW; for image acquisition, the laser was tuned to 925 nm with an intensity of 1-14 mW (both power values were measured behind the objective lens). A 60X water-immersion objective lens (Olympus) was used for both photo-labeling and image acquisition. A 40X water-immersion objective lens (Olympus) was used for image acquisition in some experiments (the ones described in Figure 2 and Figure 3). A GaAsP detector (Hamamatsu Photonics) and PMT detector were used for measuring green and red fluorescence, respectively. Photo-labeling and image acquisition files were visualized on a computer using the Prairie View software version 5.4 (Bruker). Image acquisition was performed at a resolution of 512 by 512 pixels, with a pixel size of 0.39 μm (60X lens) or 0.582 μm (40X) and a pixel dwell time of 4 μs. Each pixel was scanned 8 times.

For photo-labeling of the dorsal accessory calyx (or the posterior lateral protocerebrum), a volume spanning the entire neuropil was divided into eight to 12 planes with a step size of 2 μm. The mask function of the Prairie View software was used to mark the targeted region in every plane, and the boundaries of the mask were determined based on the red fluorescent protein DsRed expressed by the α/β_p_ Kenyon cells. The photo-labeling step was performed using a pixel size of 0.019 μm and a pixel dwell time of 2 μs. Each pixel was scanned four times. Each plane was scanned 30 times with 30-second interval. The entire photo-labeling cycle was repeated two to three times, with a 10-minute resting period between cycles. The entire brain was imaged before and after photo-labeling using the 40X water-immersion objective lens (Olympus, Japan). The number of cell bodies recovered after the photo-labeling was measured using the Multi-Point Tool function of the Fiji software (Schindelin et al., 2012).

For the photo-labeling of single neurons, a slightly modified protocol was used. Instead of a volume, a single square plane of 1.0 μm by 1.0 μm centered on the soma of a neuron cell was scanned 70 to 100 times with a 10 seconds interval between scans. Photo-labeled brains were fixed in 2% paraformaldehyde diluted in 1X phosphate bovine saline (PFA) for 45 minutes at room temperature, washed five times in 0.3% Triton X-100 diluted in 1X phosphate bovine saline (PBST) at room temperature, blocked in 5% normal goat serum diluted in PBST (PBST-NGS) for 30 minutes at room temperature, and incubated with the primary mouse antibody nc82 (1:20 in PBST-NGS; Developmental Studies Hybridoma Bank, University of Iowa) at 4°C overnight. On the following day, brains were washed four times in PBST and incubated with the secondary goat antibody Alexa Fluor 633 anti-mouse (1:500 in PBST-NGS; Life Technologies) at 4°C overnight. On the following day, brains were washed four times in PBST and mounted on a slide (Fisher Scientific) using the mounting media VECTASHIELD (Vector Laboratories Inc.). Immuno-stained brains were imaged using an LSM 880 confocal microscope. (Zeiss). The neuropils innervated by the input neurons were identified by comparing the confocal images with the adult brain template JFRC2 available on Virtual Fly Brain (https://v2.virtualflybrain.org/).

### Immunohistochemistry

For the experiments using the green fluorescent protein reconstitution across synaptic partners (GRASP) technique as well as the experiments using DenMark and synaptotagmin::GFP immunostainings, the brains of five to 14 day-old flies were dissected in saline and fixed in PFA for 45 minutes at room temperature, washed five times in PBST at room temperature, blocked in PBST-NGS for 30 minutes at room temperature, and incubated with the primary antibody at 4°C overnight. The mouse anti-GFP-20 antibody was used in GRASP and synaptotagmin experiments (1:100 in PBST-NGS; Sigma-Aldrich); the mouse nc82 antibody was used in synaptotagmin::GFP and DenMark experiments. On the following day, brains were washed four times in PBST and incubated in secondary antibody at 4°C overnight. The AlexaFluor-488 goat-anti-mouse (1:500 in PBST-NGS; Life Technologies) and AlexaFluor-633 goat-anti-mouse (1:500 in PBST-NGS; Life Technologies) were used. On the following day, brains were washed four times in PBST and mounted on a slide (Fisher Scientific) using the mounting media VECTASHIELD (Vector Laboratories Inc.). Immuno-stained brains were imaged using an LSM 880 confocal microscope. (Zeiss). The neuropils innervated by the input neurons were identified by comparing the confocal images with the adult brain template JFRC2 available on Virtual Fly Brain (https://v2.virtualflybrain.org/) (Jenett et al., 2012).

### Measuring the connectivity rate between input neurons and α/β_p_ Kenyon cells

One to three-day-old flies were used when mapping the connectivity rate between input neurons and α/β_p_ Kenyon cells. Brains were dissected in saline, treated for one minute with 2 mg/ml collagenase (Sigma-Aldrich) and mounted on a piece of Sylgard placed at the bottom of a Petri dish. The imaging protocol is the same as described above but the photo-labeling protocol is different. Each of the input neurons was photo-labeled using a single plane centered on either its soma or its projection and by scanning the plane three to five times. Each pixel was scanned eight times with a pixel size of 0.019 μm and a pixel dwell time of 4 μs. A fire-polished borosilicate glass pipette (0.5 mm I.D., 1.0 mm O.D., 10 cm length; Sutter Instruments) was pulled using the P-2000 micropipette puller (Sutter Instruments) and backfilled with Texas Red dye (lysine-fixable 3000 MW; Life Technologies) dissolved in saline. The tip of the pipette was positioned next to the cell body of a randomly chosen α/β_p_ Kenyon cell under the two-photon microscope. The dye was electroporated into the cell body using three to five 1-5 millisecond pulses of 20-50 V. The dye was allowed to diffuse within the Kenyon cell for 5 minutes before the brain was imaged.

### Confocal image acquisition and analysis

All confocal images were collected on LSM880 confocal microscope (Zeiss). For imaging whole brain, each sample was imaged twice using a Plan-Apochromat 40X/1.3 Oil M27 objective lens. First, the entire brain was divided into four tiles, each tile was imaged separately (voxel size = 0.46 μm by 0.46 μm by 2 μm, 764 by 764 pixels per image plane) and then stitched together using the Stitch function of the Zen microscope software (Zeiss). For imaging specific neuropil regions, the same objective lens was used, but with higher resolution (voxel size = 0.09 μm by 0.09 μm by 1 μm, 2320 by 2320 pixels per image plane). For imaging brains manipulated using the GRASP technique, a Plan-Apochromat 63X/1.4 Oil M27 objective lens was used in combination with the RGB Airyscan mode. Images were processed using the Airyscan function of the Zen microscope software. All confocal images were analyzed using the Fiji software (Schindelin et al., 2012). All figure panels are maximum intensity projection of confocal stacks or sub-stacks.

## AKNOWLEDGMENTS

We thank Florian Maderspacher, Kaitlyn Ellis and members of the Caron laboratory for comments on the manuscript; Shannon Torstrom for the initial characterization of the *y^1^w^1118^;;R19H07-GAL4AD^VK00027^;R20G07-GAL4DBD^attP2^*; transgenic line; Adam Lin for preparation of the standard cornmeal agar medium; Hayley Smihula and Adam Weinbrom for assistance with general laboratory concerns. This work has been funded in part by grants from the National Institute for Neurological Disorders and Stroke (R01 NS 106018 and R01 NS 1079790). Further financial support was provided by the Georges S. and Dolores Eccles Foundation (SJCC), the Dale A. Stringfellow Fellowship (JL), and the Research Scholar Award (MSJ).

## COMPETING INTERESTS

The authors declare that no competing interests exist.

## Notes

### Competing Interest Statement

The authors have declared no competing interest.

